# Connectomic dopamine-neuron disinhibition accelerates behavioral extinction

**DOI:** 10.64898/2026.08.24.746800

**Authors:** Sasha C.V. Burwell, Rene K. Carter, Haidun Yan, Shaun S.X. Lim, Brenda C. Shields, Michael R. Tadross

## Abstract

Animals must balance persistence with flexibility when outcomes change. Behavioral extinction—the reduction in responding to cues that no longer predict reward—is widely thought to be driven by pauses in dopamine-neuron firing upon omission of expected rewards. However, whether these pauses are necessary remains unknown. We used a connectomic intervention to weaken inhibitory synapses onto dopamine neurons in the ventral tegmental area, attenuating electrophysiologically defined pauses while sparing tonic and burst firing. Contrary to canonical accounts, this intervention accelerated rather than delayed behavioral extinction, demonstrating that these synapses normally postpone behavioral extinction. Learning about a newly rewarded cue was spared, arguing against a general disruption of learning. Dopamine photometry showed that the intervention attenuated reward-omission dopamine dips, whose dissipation preceded and predicted behavioral extinction across mice. These findings show that pause-generating inhibitory inputs to dopamine neurons sustain behavioral persistence. This mechanism may protect established associations from premature abandonment as outcomes fluctuate.

## Main Text

Reward-omission dopamine pauses are among the most influential neural signatures in reinforcement learning. When an expected reward fails to arrive, ventral tegmental area dopamine (VTA^DA^) neurons briefly pause their firing^1–3^, and striatal dopamine transiently falls below baseline^4^. These pauses are widely interpreted as negative prediction errors that promote associative updating and behavioral extinction^5–9^. Yet whether endogenously generated reward-omission pauses are necessary for behavioral extinction remains unresolved. Prior perturbation studies imposed dopamine-neuron excitation or inhibition at selected task events^4,8–12^; notably, excitation at reward omission replaced the endogenous pause with an opposing artificial signal^4^. These experiments established that imposed dopamine signals can influence learning but could not isolate the function of the endogenous omission pause itself. This distinction leaves open whether naturally generated reward-omission pauses serve as the required teaching signal for behavioral extinction or instead mark a circuit state with a different behavioral function.

To address this question, we attenuated dopamine pauses at their synaptic origin using gabazine^DART^, a cell-specific GABA_A_ receptor antagonist that acutely tethers to VTA^DA^ neurons^13,14^. This connectomic strategy weakens inhibitory synaptic transmission onto genetically targeted dopamine neurons, without imposing exogenously timed spike perturbations or broadly disrupting GABA release or its reception by non-dopaminergic cells. We then asked how attenuating these synapses affected dopamine-neuron dynamics and behavioral extinction.

### Gabazine^DART^ selectively attenuates GABA_A_-mediated dopamine pauses

Our approach is illustrated in spike rasters from an awake head-fixed mouse not exposed to cues or rewards, where a representative VTA^DA^ neuron exhibits tonic firing at ∼five spikes per second (black ticks), with sporadic bursts (blue) and pauses (red circles) driven by excitatory and inhibitory inputs (**Fig. 1a**, raster pre-infusion). The pauses occurred irregularly in time, making them difficult to predict or counteract using exogenously timed excitation. By contrast, rendering this neuron less sensitive to inhibitory synaptic input selectively attenuated pauses without introducing exogenous activity patterns (**Fig. 1a**, raster post-infusion).

**Fig. 1.**
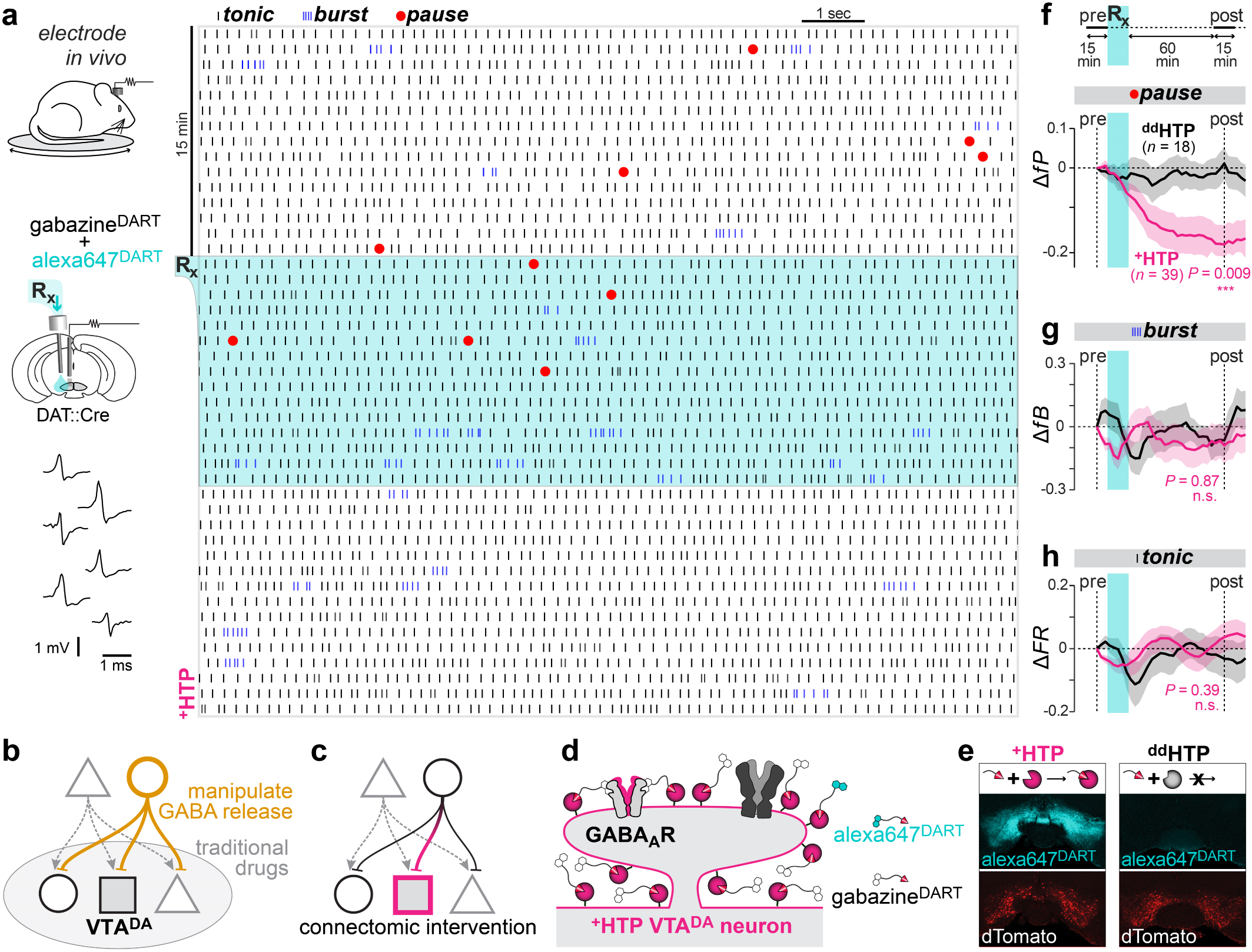
Gabazine^DART^ selectively blocks dopamine pauses. (a) Experimental setup and representative spike raster. Top left: awake head-fixed mouse on a running wheel; movable electrode targets the medial VTA and a nearby cannula permits ligand infusion. Bottom left: example extracellular spikes from putative dopamine neurons. Raster: each vertical tick represents a single spike: tonic firing (black), bursts (blue; ≥3 spikes with first and subsequent ISIs <0.08 and <0.16 sec), and pauses (red dots; ISI > 2 × the median). Each row represents 1 min (13 sec subset shown) spanning baseline (top), infusion (blue shading), and 15-min post-infusion (bottom). (b) Limitations of prior methods. Manipulations of GABA release or traditional pharmacology broadly impact both dopaminergic and non-dopaminergic cells within the VTA. (c) Connectomic strategy. GABA_A_ receptors are blocked selectively on dopamine neurons, without impacting other receptors or neighboring cell types. Acute manipulation avoids compensatory circuit adaptations. (d) DART technology. Cre-dependent AAV expression of membrane-anchored HaloTag protein (^+^HTP; pink) enables covalent capture of gabazine^DART^ (black) and alexa647^DART^ (cyan). Once tethered, gabazine^DART^ locally blocks native GABA_A_ receptors, while alexa647^DART^ enables fluorescent visualization of target engagement. (e) Representative histology. VTA_DA_ neurons expressing ^+^HTP or ^dd^HTP are labeled with dTomato (red). Alexa647^DART^ (cyan) reports ligand target engagement. (f) Pause analysis. Change in *fP* (fraction of ISI > 2 × the median) was assessed by comparing a 15-min baseline (*fP*_pre_) to a sliding 15-min window (*fP*_post_) using Δ*fP* = (*fP*_post_ *-fP*_pre_) / (*fP*_post_ + *fP*_pre_). Shading indicates mean ± SEM across cells (^dd^HTP: *n* = 18 cells, 3 mice; ^+^HTP: *n* = 39 cells, 5 mice). At 1-hr post-gabazine^DART^, Δ*fP* differed significantly between ^+^HTP and ^dd^HTP cells (*P* = 0.009, two-sided permutation test; see **fig. S2c**). (**g**-**h**) Burst and tonic analyses. *fB* (fraction of spikes fired in bursts) and *FR* (firing rate); changes quantified using similar analyses as in panel C. No significant ^+^HTP vs ^dd^HTP differences were found in these or any other non-pause metric (**fig. S2**).

We achieved this connectomic perturbation using Drug Acutely Restricted by Tethering (DART) to acutely block GABA_A_ receptors on VTA^DA^ neurons without disrupting other receptors or cell types^13,14^. By contrast, traditional GABA_A_ pharmacology^7^ and manipulations of GABA release^2,15,16^ broadly perturb GABA signaling onto dopaminergic and non-dopaminergic cells in the VTA (**Fig. 1b**), while conditional knockout of GABA_A_ receptors is confounded by chronic circuit compensation^17^. Connectomic approaches overcome these limitations with acute, intersectional precision—in this case, between genetically defined postsynaptic cells and neurotransmitter-defined presynaptic partners (**Fig. 1c**).

DART implements this perturbation in two steps. First, a Cre-dependent adeno-associated virus (AAV) drives membrane-anchored HaloTag protein (HTP) expression on the postsynaptic cell type of interest: VTA^DA^ neurons in DAT::Cre mice. Second, the perturbation is later initiated by applying gabazine^DART^, a bifunctional ligand whose HaloTag ligand (HTL) is covalently captured by HTP (**Fig. 1d**). Once tethered, the gabazine moiety is confined to a ∼5-nm shell on the VTA^DA^ surface, exerting a strong local block of GABA_A_ receptors without transcellular off-target effects^14,18^. In acute slices, ambient gabazine^DART^ leaves control dopamine neurons unaffected, but rapidly blocks GABA_A_-mediated synaptic currents in ^+^HTP-expressing dopamine neurons (**fig. S1a**)^14^. The same perturbation leaves glutamatergic transmission, intrinsic pacemaking and spike waveforms unchanged (**fig. S1b-d**)^14^.

Extracellular recordings from awake, head-fixed mice measured spontaneous single-unit spiking before and after gabazine^DART^ infusion. We identified putative dopamine neurons using established electrophysiological criteria known to yield ∼12% false-positives (**Methods**)^19–23^. Alexa647^DART^ was co-delivered to quantify drug capture histologically. Control animals were treated identically except that they expressed “double-dead” ^dd^HTP, which cannot bind HTL, thereby controlling for nonspecific effects of viral expression and ambient ligand, without inducing a tethered-drug effect (**Fig. 1e**). Histology confirmed ligand capture and specificity of viral expression: nearly all HTP expressing cells (99.7%) were dopaminergic and most dopaminergic cells in the VTA (∼64%) expressed HTP (**fig. S2, a-b**).

Tonic, burst, and pause metrics were quantified with a sliding-window analysis (**fig. S2c**). In ^dd^HTP controls, all metrics remained stable post-gabazine^DART^, indicating negligible impact of untethered ligand (**fig. S2d**). By contrast, in ^+^HTP mice, gabazine^DART^ markedly reduced spontaneous pauses, evident in the pause metric *fP* (fraction of interspike intervals exceeding twice the median), which was reduced in ^+^HTP mice relative to ^dd^HTP controls (*P* = 0.009, two-sided permutation test, **Fig. 1f**). This effect was robust to alternative pause definitions and could not be explained by symmetrical changes in interspike-interval variance (**fig. S2e**). Reductions in pause number correlated with shortening of remaining pauses (Pearson’s *r*^2^ = 0.37, *P* = 0.00004, **fig. S2f**), further confirming a pause-reduction effect. By contrast, burst and tonic-firing parameters were insensitive to tethered gabazine^DART^ (**Fig. 1, g-h** and **fig. S2c**), showing no correlations with its effects on pauses (**fig. S2g**). Thus, gabazine^DART^ attenuates GABA_A_-mediated pauses in dopamine neurons while preserving their tonic-and burst-firing characteristics.

### Gabazine^DART^ accelerates behavioral extinction without impairing new cue-reward learning

Canonical reward-prediction-error accounts propose that reward-omission pauses drive extinction^5-9^. By attenuating these pauses, gabazine^DART^ should therefore delay extinction. We tested this hypothesis in water-deprived, head-fixed mice, trained for 10 days to associate cue *A* (2.5 kHz tone, 1.5 s) with sucrose-water reward (**Fig. 2a**, top). We used cue-*A* prelicking (anticipatory licking during cue *A*, before the scheduled outcome) as the primary behavioral readout (**Fig. 2a**, left column; **Fig. 2b)**. On day 11, mice received gabazine^DART^ and brief resumption of the established *A*→reward rule, allowing measurement of cue-*A* responses under gabazine^DART^ before the contingency changed (**Fig. 2a**, “pre-flip”). *A*→omit extinction then began (**Fig. 2a**, “post-flip”) and proceeded into the final day.

**Fig. 2.**
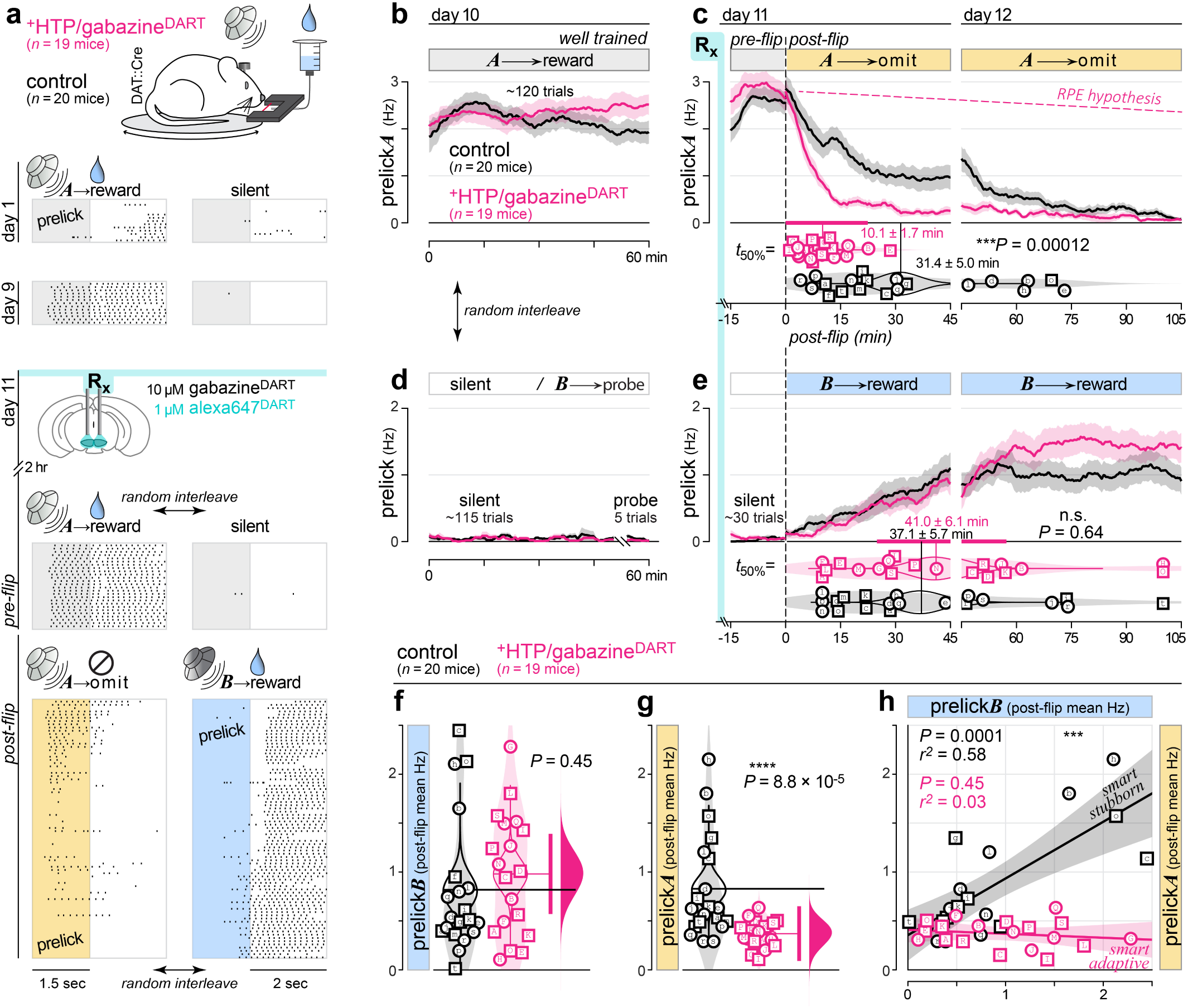
Gabazine^DART^ advances Pavlovian extinction without impairing new learning. **(a)** Pavlovian task. Mice expressing ^+^HTP or ^dd^HTP in VTA^DA^ neurons were head-fixed and trained to associate cue *A* with sucrose-water (*A*→reward). Prelicking during the cue (before reward) was measured with an infrared beam. Representative lick rasters from an ^+^HTP mouse shown for days 1, 9, and 11. On day 11, mice received an infusion of either 10 μM gabazine^DART^ + 1 μM alexa647^DART^ or 10 μM blank^DART^ + alexa647^DART^. Behavior began two hours later, starting with resumption of *A*→reward for 15 min (pre-flip), after which contingencies reversed (post-flip): *A*→omit and *B*→reward trials were randomly interleaved and continued through day 12. Throughout, *n* = 19 ^+^HTP/gabazine^DART^ and *n* = 20 control mice. **(b)** Stable cue-*A* prelicking during *A*→reward trials on day 10 (∼120 trials). Prelicking (during the 1.5 s cue) is shown as raw lick rate in Hz. Lines and shading indicate mean ± SEM. **(c)** Accelerated extinction of cue-*A* prelicking after the rule flip. **Top,** cue-A prelicking across days 11 and 12, shown as raw lick rate in Hz. Lines and shading indicate mean ± SEM. The dashed pink line illustrates the reward-prediction-error hypothesis that attenuating omission pauses should slow extinction. **Bottom,** half-extinction times (*t*_50%_) estimated from logistic fits for individual mice. Shown are individual mice (male, square; female, circle), kernel density estimates (shaded violins), bootstrap distributions of the mean (outlined violins), and group means (vertical lines). Extinction occurred approximately threefold earlier in ^+^HTP/gabazine^DART^ mice (mean ± SEM, 10.1 ± 1.7 min) than in controls (31.4 ± 5.0 min). The thick horizontal pink bar indicates the 95% confidence interval from a two-sided permutation test, showing a significant difference between treatment groups (*P* = 0.00012). Lettered symbols identify the same mice throughout the manuscript. **(d)** Cue-discrimination control. On day 10, prelicking remained near zero during silent trials (∼115 trials) and five unrewarded cue-*B* probe trials. The probe trials were interspersed throughout the 60-min session but are plotted together at the end for display. **(e)** Intact new cue–reward learning. Prelicking remained near zero during pre-flip silent trials (∼30 trials) and emerged during post-flip *B*→reward conditioning. Half-conditioning times did not differ detectably between ^+^HTP/gabazine^DART^ (41.0 ± 6.1 min) and control (37.1 ± 5.7 min) mice (*P* = 0.64, two-sided permutation test). Format as in panel **c**. **(f)** Mean cue-*B* prelicking across the post-flip period. Shown are individual mice (male, square; female, circle), kernel density estimates (shaded violins), bootstrap distributions of the group means (outlined violins), and group means (horizontal lines). At right, hypothesis testing is shown as the 95% CI from a two-sided permutation test (vertical pink bar) and effect size as the bootstrap distribution of the mean difference (pink half-violin). Cue-*B* prelicking did not differ detectably between ^+^HTP/gabazine^DART^ and control mice (*P* = 0.45, two-sided permutation test). **(g)** Mean cue-*A* prelicking across the post-flip period, displayed as in **f**. Cue-*A* prelicking was lower in ^+^HTP/gabazine^DART^ than in control mice, indicating accelerated extinction (*P* = 8.8 × 10⁻⁵, two-sided permutation test). **(h)** Within-mouse relationship between mean post-flip cue-*B* and cue-*A* prelicking. Symbols denote individual mice (male, square; female, circle). In control mice, greater cue-*B* prelicking was associated with more persistent cue-*A* prelicking—the “smart-stubborn” relationship (Pearson’s *r*² = 0.58, *P* = 0.0001). This relationship was not detectable in ^+^HTP/gabazine^DART^ mice, which instead showed uniformly low cue-*A* prelicking despite variable cue-B conditioning (*r*² = 0.03, *P* = 0.45). The relationships differed significantly between groups (treatment × cue-*B* prelicking interaction, *P* = 6.8 × 10⁻⁵). Lines and shading indicate linear fits and 95% confidence intervals.

Gabazine^DART^ did not alter cue-*A* prelicking during the initial *A*→reward block (pre-flip mean: ^+^HTP/gabazine^DART^ = 2.82 ± 0.24 Hz, *n* = 19 mice; controls = 2.49 ± 0.19 Hz, *n* = 20 mice; *P* = 0.28, two-sided permutation, **fig. S3a-b**), arguing against a broad change in motivation or response vigor. By contrast, cue-*A* prelicking during *A*→omit trials reached half-extinction approximately threefold earlier in gabazine^DART^-treated mice (*t*_50%_ = 10.1 ± 1.7 min, mean ± SEM, *n* = 19 mice) than in controls (*t*_50%_ = 31.4 ± 5.0 min, *n* = 20 mice; *P* = 0.00012, two-sided permutation)—an effect opposite to the prediction that reward-omission pauses are required for extinction (**Fig. 2c**).

To ensure that extinction of cue-*A* prelicking reflected associative learning, we used a design that maintained a constant overall reward rate while reassigning reward to a novel cue *B* (11 kHz tone, 1.5 s). This approach has been shown to mitigate nonspecific effects of global reward loss, allowing declining cue-*A* prelicking to be interpreted as cue–outcome learning rather than nonspecific motivational reduction^24^. The protocol also combined training and quality control to ensure that mice treated cue *A* and cue *B* as distinct stimuli. Throughout the assay, licking during inter-trial intervals triggered timeouts; this trained mice to withhold licking to non-cue-*A* events during days 1–9, when cue *B* was withheld to preserve its novelty and *A*→reward trials were interleaved with silent trials (**Fig. 2a**, right column; **fig. S3c**). As a quality-control test on day 10, five silent trials were replaced by unrewarded cue-*B* probes; 39 of 41 mice that learned to prelick to cue *A* did not prelick to cue *B*, even on its first presentation, and the two that failed this test were excluded from further analysis (**Fig. 2d**, probe trials).

Interleaved *B*→reward trials also provided a within-mouse measure of new Pavlovian conditioning (**Fig. 2e**), which was unaffected by gabazine^DART^ (*t*_50%_ = 41.0 ± 6.1 min, *n* = 19 mice) relative to controls (*t*_50%_ = 37.1 ± 5.7 min, *n* = 20 mice; *P* = 0.64, two-sided permutation). Thus, new cue-reward learning remained intact in the same animals where gabazine^DART^ advanced the onset of extinction.

Individual control mice varied in both cue-*B* acquisition (**Fig. 2f**, black) and extinction resistance, measured as persistent cue-*A* prelicking (**Fig. 2g**, black). Across the same mice, these measures were positively correlated: mice with greater cue-*B* prelicking during conditioning also showed more persistent cue-*A* prelicking during extinction (*P* = 0.0001, *r*² = 0.58, *n* = 20; **Fig. 2h**, black). We refer to this continuous relationship as the “smart-stubborn” correlation; mice in the upper right expressed both traits most strongly.

Gabazine^DART^-treated mice spanned the full range of cue-*B* acquisition as in controls (prelick*B*, post-flip mean: gabazine^DART^, 0.98 ± 0.14 Hz, *n* = 19 mice; controls, 0.82 ± 0.16 Hz, *n* = 20 mice; *P* = 0.45, two-sided permutation, **Fig. 2f**) but showed consistently little persistent cue-*A* prelicking (prelick*A*, post-flip mean: gabazine^DART^, 0.37 ± 0.03 Hz, *n* = 19 mice; controls, 0.83 ± 0.12 Hz, *n* = 20 mice; *P* = 8.8 × 10^-5^, two-sided permutation, **Fig. 2g**). Consequently, a smart-stubborn correlation was not detectable in gabazine^DART^-treated mice (*P* = 0.45, *r*² = 0.03, *n* = 19; **Fig. 2h**, pink), and a treatment × cue-*B*-prelicking interaction confirmed that the relationship differed between groups (*P* = 6.8 × 10^-5^). Thus, gabazine^DART^ decoupled new learning from extinction resistance, yielding “smart-adaptive” individuals that both acquired and extinguished effectively—a phenotype not seen in controls (**Fig. 2h**, bottom right). Taken together, gabazine^DART^ advanced the onset of extinction without compromising new cue–reward learning.

### Distinct dopamine dynamics dissociate behavioral extinction from locomotor activation

In a subset of mice, we concurrently monitored extracellular dopamine dynamics during the Pavlovian assay with GRAB_DA3m_ photometry in the nucleus accumbens^25^. We predefined seven behavioral epochs: the inter-trial interval (ITI), pre-flip cue-*A* presentation (preflipCue*A*) and associated reward (reward*A*), post-flip cue-*A* presentation (cue*A*) and associated omission (omit), and post-flip cue-*B* presentation (cue*B*) and associated reward (reward*B*). Cue windows were aligned to the 1.5 s cues, whereas outcome windows spanned the 2 s following cue offset.

Because fiber photometry cannot resolve electrophysiologically defined pauses and bursts, we separated the continuous fluorescence-derived dopamine signal into downward-going and upward-going components by half-wave rectification (**fig. S4a**, steps 1-2). We sign-inverted the downward-going component and denoted the resulting waveforms DA↓(t) and DA↑(t); both waveforms were non-negative whereas the arrows indicate the original direction of deflection (**fig. S4a**, step 2). We then isolated each of the seven predefined behavioral epochs and visualized how each signal evolved across the assay on a common minutes-scale axis (**fig. S4a**, step 3). For example, DA↓^ITI^(t) shows how the magnitude of downward-going deflections during ITIs evolved across the assay, revealing a small but abrupt reduction following gabazine^DART^ infusion (**fig. S4b**, left). Finally, we used the normalized within-mouse change metric Δ = (post − pre) / (post + pre) for statistical analyses, where pre and post denote each mouse’s mean pre-and post-infusion values for the corresponding direction and epoch (**fig. S4a**, step 4). For example, positive values of ΔDA↓^ITI^ indicate deeper or more prevalent dopamine dips following treatment, whereas negative values indicate shallower or less prevalent dopamine dips (**fig. S4b**, right).

Across predefined task epochs, gabazine^DART^ preferentially attenuated dopamine dips. We therefore controlled familywise error across the seven predefined downward-going metrics (**fig. S4b-e**). Two survived Holm correction (**Fig. 3a-b**): ITI dopamine dips remained largely unchanged in controls and were attenuated by gabazine^DART^ (ΔDA↓^ITI^: controls, −0.03 ± 0.03; gabazine^DART^, −0.18 ± 0.02; *P*_unadj_ = 0.0026, *P*_Holm_ = 0.018); omission-related dopamine dips dissipated over the assay in controls and were further attenuated by gabazine^DART^ (ΔDA↓^omit^: controls, −0.52 ± 0.03; gabazine^DART^, −0.66 ± 0.03; *P*_unadj_ = 0.0077, *P*_Holm_ = 0.046). Other downward-going metrics showed directionally similar effects that did not survive Holm correction (**fig. S4b-e**). A parallel analysis of the seven upward-going metrics revealed no nominally significant or Holm-corrected effects of gabazine^DART^ (all *P*_unadj_ ≥ 0.077, *P*_Holm_ ≥ 0.54; **fig. S5**). Accordingly, subsequent behavioral–photometry analyses focused on ΔDA↓^ITI^ and ΔDA↓^omit^.

**Fig. 3.**
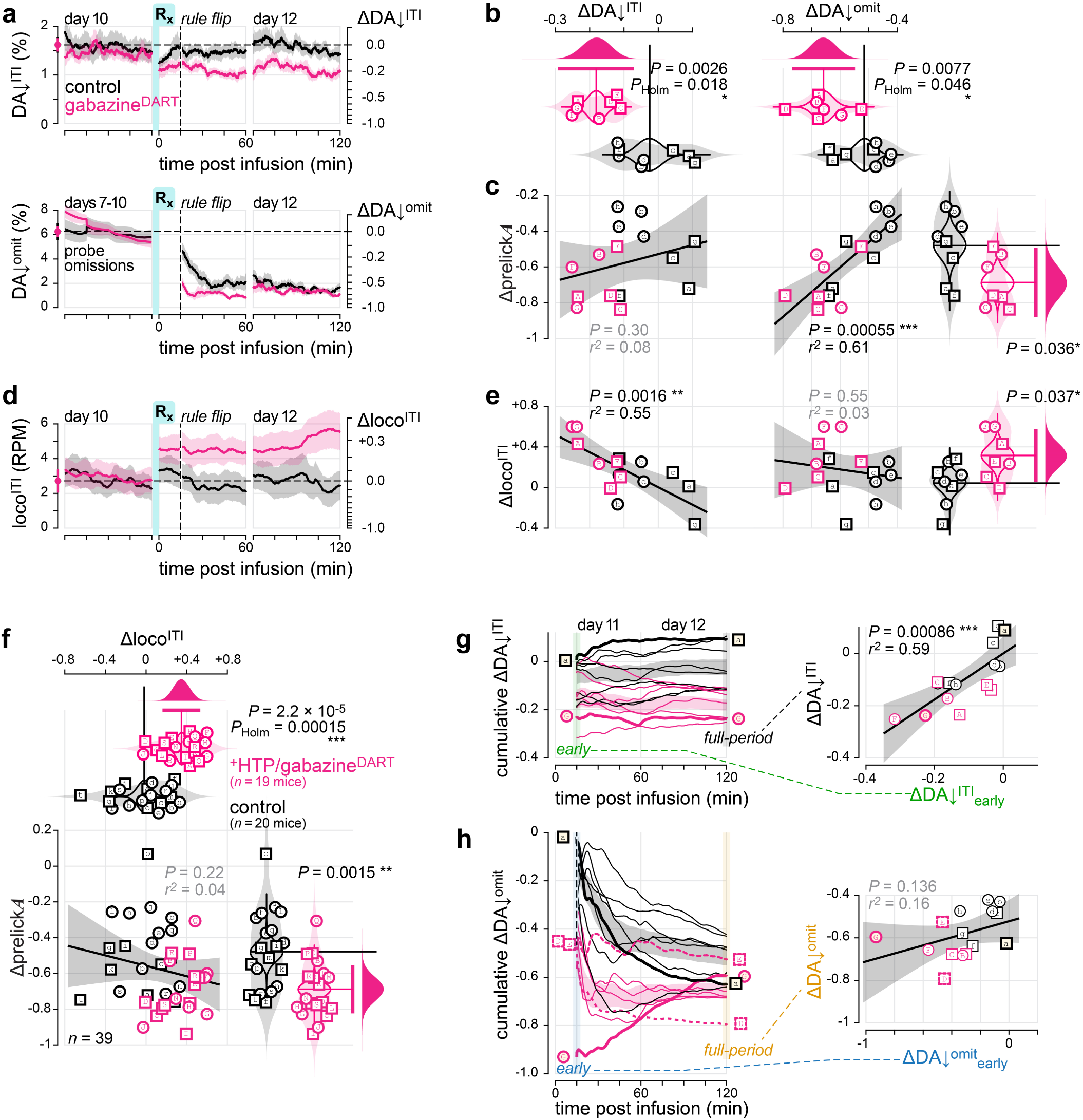
Distinct dopamine dynamics dissociate behavioral extinction from locomotor activation. **(a)** Time courses of the two downward-going nucleus accumbens dopamine signals that survived Holm correction for family-wise error. See **fig. S4a** for the rectified epoch-analysis workflow. For display only, data from each treatment group were multiplied by a single constant so that the day 7–10 pre-infusion time average for that group equaled the geometric mean of the two group baselines (far-left symbol and dashed horizontal line). This group-level rescaling preserved relative within-group variability without normalizing individual mice. Raw values are provided in supplementary data. The left axis shows baseline-aligned signal amplitude in native percentage fluorescence-change units. The right axis shows Δ = (post − pre) / (post + pre), comparing the post-infusion waveform to the pre-infusion mean from days 7–10. **Top,** DA↓^ITI^, downward-going dopamine signals during intertrial intervals, shown across days 10–12. **Bottom,** DA↓^omit^, downward-going dopamine signals during reward omission. For DA↓^omit^, pre-infusion values were measured during probe omissions on training days 7–10 (three per day). Lines and shading indicate mean ± s.e.m. Pink, ^+^HTP/gabazine^DART^ (*n* = 7 mice); black, control (*n* = 8 mice). **(b)** Gabazine^DART^ attenuated both DA↓^ITI^ and DA↓^omit^ across the full post-infusion period. More negative ΔDA↓^ITI^ and DA↓^omit^ values indicate greater dissipation of downward-going dopamine signals following drug infusion. Shown are individual mice (male, square; female, circle), kernel density estimates (shaded violins), bootstrap distributions of the group means (outlined violins), and group means (vertical lines). The 95% CI from a two-sided permutation test is shown as a horizontal pink bar, and the bootstrap distribution of the mean difference is shown as a pink half-violin. Gabazine^DART^ attenuated both DA↓^ITI^ (*P*_unadj_ = 0.0026; *P*_Holm_ = 0.018) and DA↓^omit^ (*P*_unadj_ = 0.0077; *P*_Holm_ = 0.046). Holm correction was applied across seven predefined behavioral epochs (**fig. S4b-e**). **(c)** Behavioral extinction tracked DA↓^omit^ but not DA↓^ITI^. **Left,** Δprelick*A* was not detectably associated with ΔDA↓^ITI^ (Pearson’s *P* = 0.30, *r*² = 0.08). **Middle,** Δprelick*A* was strongly associated with ΔDA↓^omit^ (*P* = 0.00055, *r*² = 0.61). **Right,** full-period change in cue-*A* prelicking. More negative Δprelick*A* values indicate more efficient extinction of cue-*A* prelicking. Gabazine^DART^ enhanced behavioral extinction relative to control treatment (*P* = 0.036, two-sided permutation test), displayed as in **b**. **(d)** Gabazine^DART^ increased locomotion during intertrial intervals. The time course of loco^ITI^ (wheel rotations per min) across days 10–12 is displayed as in **a**. The left axis shows baseline-aligned locomotion in rotations per min. The right axis shows the normalized within-mouse change, Δloco^ITI^. Lines and shading indicate mean ± s.e.m. Pink, ^+^HTP/gabazine^DART^ (*n* = 19 mice); black, control (*n* = 20 mice). See **fig. S6** for locomotor analyses across other behavioral epochs. **(e)** Locomotor activation tracked DA↓^ITI^ but not DA↓^omit^. **Left,** Δloco^ITI^ was associated with ΔDA↓^ITI^ (Pearson’s *P* = 0.0016, *r*² = 0.55). **Middle,** Δloco^ITI^ was not detectably associated with ΔDA↓^omit^ (*P* = 0.55, *r*² = 0.03). **Right,** gabazine^DART^ increased ITI locomotion relative to control treatment within the photometry subset of mice (*P* = 0.037, two-sided permutation test; pink, ^+^HTP/gabazine^DART^, *n* = 7 mice; black, control, *n* = 8 mice), displayed as in **b**. **(f)** Locomotor activation and behavioral extinction were not detectably associated in the full behavioral cohort (*n* = 39 mice; 19 ^+^HTP/gabazine^DART^ and 20 controls). **Top,** gabazine^DART^ increased ITI locomotion (*P*_unadj_ = 2.2 × 10⁻⁵; *P*_Holm_ = 0.00015), displayed as in **b**. **Bottom left,** Δloco^ITI^ was not detectably associated with Δprelick*A* across mice (Pearson’s *P* = 0.22, *r*² = 0.04). **Bottom right,** gabazine^DART^ produced greater behavioral extinction than control treatment (*P* = 0.0015, two-sided permutation test), displayed as in **b**. See **fig. S7a** for the corresponding comparison of ΔDA↓^ITI^ and ΔDA↓^omit^. **(g)** Early attenuation of DA↓^ITI^ remained stable across the full post-infusion period. ΔDA↓^ITI^ was calculated cumulatively: at each plotted time *t*, the fixed pre-infusion mean was compared with the mean post-infusion signal from 0 to *t*. Thus, the first point, ΔDA↓^ITI^_early_, is plotted at 15 min and used the post-infusion mean from 0–15 min; the point at 60 min uses the mean from 0–60 min; and the full-period ΔDA↓^ITI^ is plotted at 120 min and uses the mean from 0–120 min. **Left,** cumulative trajectories for individual mice. Shading indicates mean ± s.e.m. Pink, ^+^HTP/gabazine^DART^ (*n* = 7 mice); black, control (*n* = 8 mice). **Right,** relationship between ΔDA↓^ITI^_early_ and full-period ΔDA↓^ITI^ (Pearson’s *P* = 0.00086, *r*² = 0.59). Lines and shading indicate the linear fit and 95% confidence interval. Lettered symbols identify the same mice throughout the manuscript. **(h)** Early attenuation of DA↓^omit^ did not predict its full-period attenuation. ΔDA↓^omit^ was calculated cumulatively as in **g**. Here, because omission trials begin at 15 min, ΔDA↓^omit^_early_ is plotted at 15 min and compares the pre-infusion mean with the mean from the first four post-flip omissions. Full-period ΔDA↓^omit^ is plotted at 120 min and uses the mean over all post-flip omissions. **Left,** cumulative trajectories for individual control (black) and ^+^HTP/gabazine^DART^ (pink) mice. Right, relationship between early and full-period ΔDA↓^omit^ (Pearson’s *P* = 0.136, *r*² = 0.16). Selected trajectories are highlighted and linked to their corresponding points. See **fig. S7b–e** for related correlation analyses.

We next asked whether either photometry measure correlated with behavioral extinction (**Fig. 3c**). Across mice, ΔDA↓^ITI^ was not detectably correlated with Δprelick*A*, the behavioral readout of extinction (*P* = 0.30, *r*² = 0.08). By contrast, ΔDA↓^omit^ was strongly correlated with Δprelick*A* (*P* = 0.00055, *r*²=0.61). Thus, downward-going dopamine dynamics during reward omissions—but not during ITIs—correlated with behavioral extinction (**Fig. 3c**).

Gabazine^DART^ also enhanced locomotion during ITIs (Δloco^ITI^: controls, −0.02 ± 0.06; gabazine^DART^, +0.35 ± 0.05; *P*_unadj_ = 2.2 × 10⁻⁵, *P*_Holm_ = 0.00015, two-sided permutation test; Fig. 3d and **fig. S6b**), with other epochs showing directionally similar but weaker effects (**fig. S6**). Critically, locomotion showed the complementary pattern of photometry correlations (**Fig. 3e**): here, ΔDA↓^ITI^ strongly correlated with locomotion (*P* = 0.0016, *r*² = 0.55), whereas ΔDA↓^omit^ was not detectably correlated (*P* = 0.55, *r*² = 0.03).

Consistent with this separation, the two behavioral measures—locomotion and prelick*A* suppression—were not detectably correlated across the full 39-animal behavioral cohort (*P* = 0.22, *r*² = 0.04, *n* = 39), even though gabazine^DART^ altered both behaviors (**Fig. 3f**). The two photometry metrics were likewise not detectably correlated across mice (ΔDA↓^ITI^ versus ΔDA↓^omit^: *P* = 0.21, *r*² = 0.12, *n* = 15; **fig. S7a**).

Multivariable models further supported this dissociation. For locomotion, ΔDA↓^ITI^ was the only significant predictor when modeled with treatment (ΔDA↓^ITI^, *P* = 0.024; treatment, *P* = 0.95) or together with ΔDA↓^omit^ (ΔDA↓^ITI^, *P* = 0.047; ΔDA↓^omit^, *P* = 0.54; treatment, *P* = 0.66). Extinction showed the opposite structure: ΔDA↓^omit^ was the only significant predictor when modeled with treatment (ΔDA↓^omit^, *P* = 0.0089; treatment, *P* = 0.85) or together with ΔDA↓^ITI^ (ΔDA↓^omit^, *P* = 0.016; ΔDA↓^ITI^, *P* = 0.98; treatment, *P* = 0.88). These analyses revealed a double dissociation: ΔDA↓^ITI^ attenuation predicted locomotor enhancement, whereas ΔDA↓^omit^ attenuation predicted behavioral extinction.

### ITI and omission dynamics diverge over time

We next asked how two downward-going dopamine measures affected by the same perturbation could become uncoupled. We reasoned that the perturbation itself should persist throughout the session, whereas omission-locked responses could evolve as reward expectations changed. We therefore compared early and full-period measures to examine their evolution over time.

An early ITI measure, ΔDA↓^ITI^early, calculated from the first 15 min of the post-infusion waveform, was strongly correlated with full-period ΔDA↓^ITI^ using all 120 min of the post-infusion waveform (*P* = 0.00086, *r*² = 0.59; **Fig. 3g**), indicating that the impact of gabazine^DART^ on DA↓^ITI^ varied across mice yet remained stable over time. We next examined an early omission measure, ΔDA↓^ITI^early, calculated from the first four post-flip omissions, which notably correlated with both ΔDA↓^ITI^early(*P* = 0.019, *r*² = 0.35) and full-period ΔDA↓ (*P* = 0.0079, *r*² = 0.43), indicating a shared acute effect of gabazine^DART^ during ITIs and the first few omissions (**fig. S7b-c**).

By contrast, full-period ΔDA↓^omit^ diverged from the other measures: it was not detectably correlated with its early counterpart, ΔDA↓^omit^ (*P* = 0.136, *r*² = 0.16; **Fig. 3h**) or with either ΔDA↓^ITI^ metric (**fig. S7d-e**). Individual time courses illustrate this divergence. The mouse with the most negative ΔDA↓^omit^ value—and thus the strongest initial gabazine^DART^ effect—showed a later rebound, yielding only an intermediate cumulative ΔDA↓^omit^ (**Fig. 3h**, mouse “G”). A similar cumulative ΔDA↓^omit^ occurred in a control despite receiving no perturbation (mouse “a”). Thus, animals with markedly different early omission dips could converge on similar cumulative values. Likewise, mice with intermediate early values could diverge (for example, mice “D” and “E”). ΔDA↓^omit^ therefore reflected both the initial effect of gabazine^DART^ and the subsequent evolution of each animal’s omission responses during repeated reward omission.

In sum, gabazine^DART^ affected ΔDA↓^ITI^ early and ΔDA↓^omit^ early in parallel. ΔDA↓^ITI^ remained stable over time, whereas ΔDA↓^omit^ evolved with repeated reward omission, resulting in a distinct omission-locked emergent signal associated with behavioral extinction.

### Behavioral extinction dissociates from cue-evoked dopamine decline

Neutral cues generally evoke little dopamine response but can come to evoke robust dopamine elevations as cues become predictive of reward. These acquired responses are widely interpreted as dopamine signatures of learned cue value^1,6^—an internal signal that typically goes hand-in-hand with conditioned responding to the same cue. Under a unitary value-updating account, the two should therefore change together as the association is acquired or extinguished. Consistent with this interpretation, DA↑^cue*A*^ was small on the first training day but grew substantially during *A*→reward training, paralleling the development of cue-*A* prelicking (**fig. S8a-b**). Across mice, DA↑^cue*A*^ and cue-*A* prelicking were positively correlated during training days 1–4 (*P* = 0.0088, *r*² = 0.42, *n* = 15 mice; **fig. S8c**). Thus, DA↑^cue*A*^ and cue-*A* prelicking both tracked the development of the learned *A*→reward association.

We therefore expected that the two readouts would also decline together during extinction. Contrary to this expectation, gabazine^DART^ separated their trajectories. Restricting analysis to the photometry cohort confirmed that gabazine^DART^ advanced behavioral extinction, both by half-time (controls: *t*_50%_ = 41.8 ± 8.6 min, *n* = 8 mice; gabazine^DART^: 11.8 ± 3.9 min, *n* = 7 mice; *P* = 0.012, two-sided permutation test; **Fig. 4a**) and normalized change (Δprelick*A*, *P* = 0.036; **Fig. 3c**). By contrast, gabazine^DART^ did not detectably alter DA↑^cue*A*^ decline, either by half-time (controls: *t*_50%_ = 45.2 ± 7.0 min; gabazine^DART^: 48.9 ± 8.6 min; *P* = 0.74; **Fig. 4b**) or normalized change (ΔDA↑^cue*A*^, *P* = 0.58; **fig. S5d**).

**Fig. 4.**
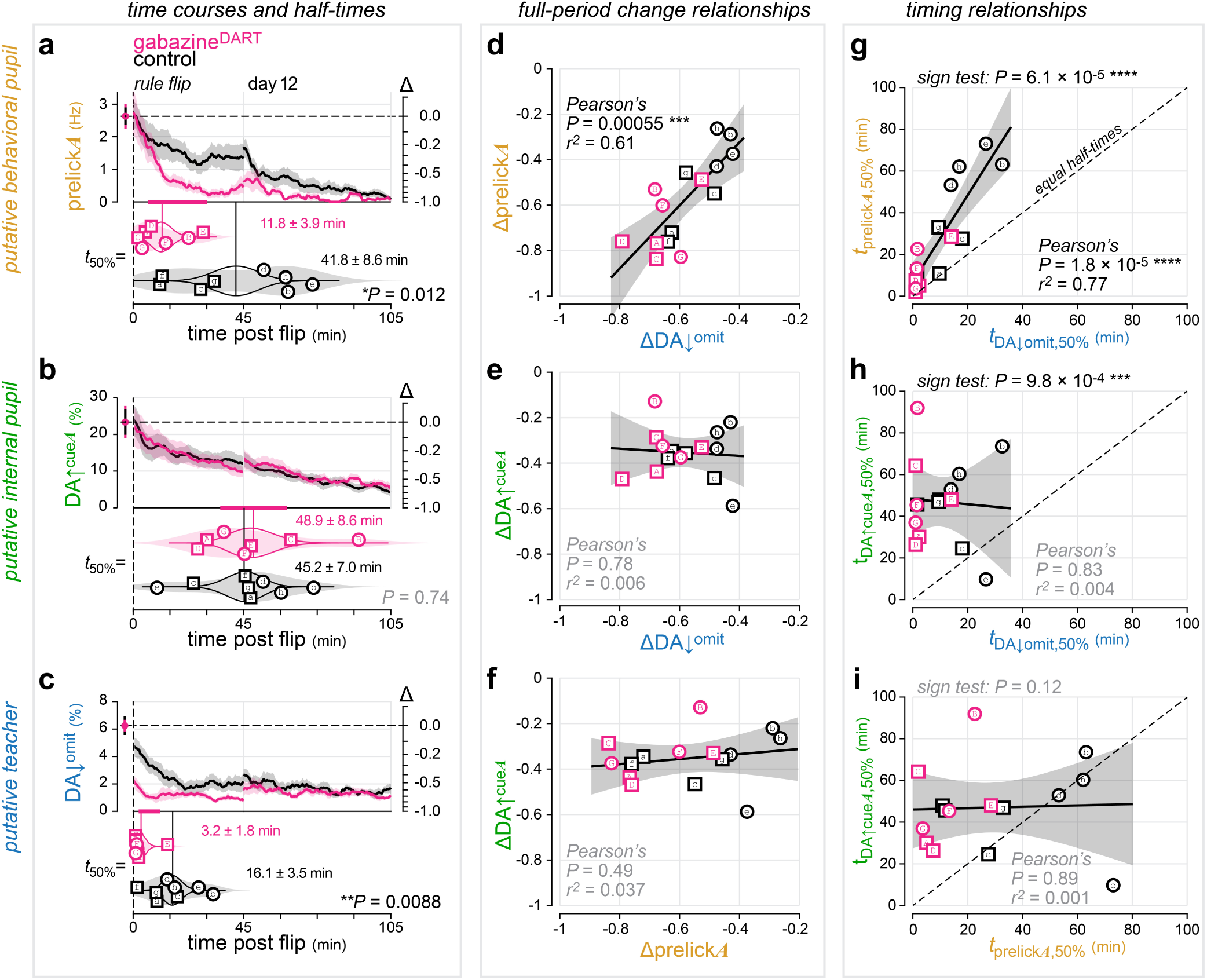
Behavioral extinction and cue-evoked dopamine decline dissociate, yet both occur after DA↓^omit^ dissipation. **(a)** Gabazine^DART^ accelerated behavioral extinction within the photometry cohort. **Top,** cue-*A* prelicking across the post-flip period on days 11 and 12. The far-left symbols and dashed horizontal line show the baseline-aligned pre-flip mean; the right axis shows normalized change relative to this value. Lines and shading indicate mean ± s.e.m. **Bottom,** half-extinction times (*t*_50%_) estimated from logistic fits for individual mice. Shown are individual mice (male, square; female, circle), kernel density estimates (shaded violins), bootstrap distributions of the group means (outlined violins), and group means (vertical lines). Hypothesis testing is shown as the 95% CI from a two-sided permutation test (horizontal pink bar). Extinction occurred earlier in ^+^HTP/gabazine^DART^ mice (11.8 ± 3.9 min) than in controls (41.8 ± 8.6 min; *P* = 0.012). **(b)** Gabazine^DART^ did not detectably alter DA↑^cue*A*^ decline. **Top,** DA↑^cue*A*^ across the post-flip period, displayed as in **a**. **Bottom,** half-decline times estimated from logistic fits, displayed as in **a**. Half-times did not differ detectably between ^+^HTP/gabazine^DART^ mice (48.9 ± 8.6 min) and controls (45.2 ± 7.0 min; *P* = 0.74, two-sided permutation test). **(c)** Gabazine^DART^ advanced dissipation of DA↓^omit^. **Top,** DA↓^omit^ across the post-flip period, displayed as in **a**. **Bottom,** half-dissipation times estimated from logistic fits, displayed as in **a**. DA↓^omit^ dissipated earlier in ^+^HTP/gabazine^DART^ mice (3.2 ± 1.8 min) than in controls (16.1 ± 3.5 min; *P* = 0.0088, two-sided permutation test). **(d)** Full-period DA↓^omit^ attenuation tracked behavioral extinction. Normalized change in cue-*A* prelicking is plotted against normalized change in DA↓^omit^ (Pearson’s *P* = 0.00055, r² = 0.61). Lettered symbols identify the same mice across panels; *n* = 7 ^+^HTP/gabazine^DART^ (pink) and *n* = 8 controls (black). Solid lines and shading indicate linear fits and 95% confidence intervals. **(e)** Full-period DA↓^omit^ attenuation did not track DA↑^cue*A*^ decline. Normalized change in DA↑^cue*A*^ is plotted against normalized change in DA↓^omit^ (Pearson’s *P* = 0.78, *r*² = 0.006). Format as in **d.(f)** Behavioral extinction and cue-evoked dopamine decline were not detectably associated. Normalized change in DA↑^cue*A*^ is plotted against normalized change in cue-*A* prelicking (Pearson’s *P* = 0.49, *r*² = 0.037). Format as ind. **(g)** Behavioral extinction occurred after DA↓^omit^ dissipation in every mouse, and their half-times tightly covaried across mice. The half-extinction time of cue-*A* prelicking is plotted against the half-dissipation time of DA↓^omit^. All 15 mice fell above the dashed line of equal half-times (two-sided sign test, *P* = 6.1 × 10⁻⁵). Half-times were positively associated across mice (Pearson’s *P* = 1.8 × 10⁻⁵, *r*² = 0.77). The regression slope was also significantly greater than 1 (*P* = 0.0048, two-sided test against slope = 1), indicating that the interval between DA↓^omit^ dissipation and behavioral extinction increased as DA↓^omit^ dissipation occurred later. In a multivariable model, DA↓^omit^ timing remained associated with behavioral extinction timing after accounting for treatment (*P* = 0.00086), whereas treatment explained no detectable additional variance (*P* = 0.50; model *r*² = 0.78, adjusted *r*² = 0.74). The treatment × DA↓^omit^ half-time interaction was also not significant (*P* = 0.76), providing no detectable evidence that the relationship differed between groups. **(h)** DA↑^cue*A*^ decline occurred after DA↓^omit^ dissipation, but their half-times did not covary. The half-decline time of DA↑^cue*A*^ is plotted against the half-dissipation time of DA↓^omit^. DA↑^cue*A*^ decline occurred after DA↓^omit^ dissipated in 14 of 15 mice (two-sided sign test, *P* = 9.8 × 10⁻⁴), whereas the two half-times were not detectably associated across mice (Pearson’s *P* = 0.83, *r*² = 0.004). **(i)** Behavioral extinction and cue-evoked dopamine decline showed neither consistent temporal ordering nor detectable half-time covariance. The half-decline time of DA↑^cue*A*^ is plotted against that of cue-*A* prelicking (two-sided sign test, *P* = 0.12; Pearson’s *P* = 0.89, *r*² = 0.001).

Gabazine^DART^ accelerated dissipation of DA↓^omit^ (controls: *t*_50%_ = 16.1 ± 3.5 min; gabazine^DART^: 3.2 ± 1.8 min; *P* = 0.0088; **Fig. 4c**), and ΔDA↓^omit^ was strongly correlated with behavioral extinction across mice (*P* = 0.00055, *r*² = 0.61; **Fig. 4d**). Yet ΔDA↓^omit^ was not detectably correlated with DA↑^cue*A*^ decline (*P* = 0.78, *r*² = 0.006; **Fig. 4e**), which was itself not detectably correlated with behavioral extinction (*P* = 0.49, *r*² = 0.037; **Fig. 4f**). Finally, gabazine^DART^ did not detectably alter the half-times of cue-*B* prelicking, DA↑^cue*B*^ emergence, or DA↑^reward*B*^ persistence (all *P* ≥ 0.15; **fig. S9a-c**). Thus, gabazine^DART^ advanced the onset of behavioral extinction without detectably accelerating the decline of the internal representation of cue value—an unexpected finding that we revisit in the Discussion.

### Omission-related dopamine dynamics dissipate before extinction emerges

We next examined the temporal relationship between each outcome-associated dopamine signal (putative teacher; Fig. 4c and **fig. S9c**) and its corresponding behavioral and cue-evoked dopamine readouts (putative behavioral and internal pupils, respectively; **Fig. 4a-b** and **fig. S9a-b**).

During *B*→reward conditioning, the outcome-associated DA↑^rewardB^ persisted beyond the emergence of cue-*B* prelicking in 14 of 15 mice (two-sided sign test, *P* = 9.8 × 10^-4^; **fig. S9d**) and beyond DA↑^cue*B*^ emergence in all 15 mice (two-sided sign test, *P* = 6.1 × 10⁻⁵; **fig. S9e**), while DA↑^cue*B*^ and cue-*B* prelicking showed no consistent ordering (two-sided sign test, *P* = 0.09; **fig. S9f**). Thus, the outcome-associated DA↑^reward*B*^ signal was consistent with a teaching signal that persists until learning has stabilized.

The cue-*A* extinction pathway showed the opposite temporal structure. The outcome-associated DA↓^omit^ dissipated before behavioral extinction in all 15 mice (two-sided sign test, *P* = 6.1 × 10⁻⁵; **Fig. 4g**) and before DA↑^cue*A*^ decline in 14 of 15 mice (*P* = 9.8 × 10⁻⁴; **Fig. 4h**). Within the gabazine^DART^ group, behavioral extinction preceded DA↑^cueA^ decline in all seven mice, whereas controls showed no consistent ordering (**Fig. 4i**). Direct within-mouse comparisons of outcome-to-readout lags confirmed the reversal between conditioning and extinction: relative to both behavioral and cue-evoked dopamine readouts, DA↑^reward*B*^ persisted later during cue-*B* conditioning than DA↓^omit^ did during cue-*A* extinction (both *P* = 6.1 × 10⁻⁵, two-sided sign tests). Thus, the outcome-associated DA↓^omit^ signal was not consistent with a teaching signal that persists as extinction emerges.

### Gabazine^DART^ preserves the relationship between DA↓^omit^ and behavioral extinction

Among the six behavioral and dopamine transitions quantified by half-time (prelick*A*, DA↑^cue*A*^, DA↓^omit^, prelick*B*, DA↑^cue*B*^ and DA↑^reward*B*^), gabazine^DART^ shifted two events earlier: DA↓^omit^ dissipation and prelick*A* extinction. This coordinated shift raised two possibilities. Gabazine^DART^ could act separately on DA↓^omit^ dissipation and behavioral change, or primarily on an upstream transition marked by DA↓^omit^ dissipation. Under the latter model, DA↓^omit^ timing should predict extinction onset after accounting for treatment, and this relationship should not differ between groups.

To distinguish these possibilities, we modeled behavioral-extinction onset as a function of DA↓^omit^ dissipation timing and treatment assignment. Across mice, DA↓^omit^ timing strongly predicted behavioral-extinction timing (*P* = 1.8 × 10⁻⁵, *r*² = 0.77; **Fig. 4g**). This relationship remained significant in a model including treatment (*P* = 0.00086), whereas treatment explained no additional variance (*P* = 0.50; model *r*² = 0.78, adjusted *r*² = 0.74). The treatment × DA↓^omit^ timing interaction was also not significant (*P* = 0.76), providing no detectable evidence that the relationship differed between groups. Thus, gabazine^DART^ shifted DA↓^omit^ dissipation earlier, with behavioral extinction following according to the same relative timing observed in controls; the data did not support an additional treatment-specific relationship with extinction (**Fig. 4g**).

Finally, we tested a delayed-priming alternative in which omission signals initiate extinction but disappear before its behavioral expression. If omission dips primed subsequent extinction, then longer omission-dip exposure should predict a shorter delay between DA↓^omit^ dissipation and behavioral extinction, yielding a slope below 1 in **Fig. 4g**. Instead, the regression slope was significantly greater than 1 (*P* = 0.0048, two-sided test against slope = 1), indicating that longer omission-dip exposure predicted a longer delay from DA↓^omit^ dissipation to behavioral extinction. Moreover, in five of seven gabazine^DART^-treated mice, DA↓^omit^ dissipated in under 2 min, leaving little opportunity for omission dips to prime extinction, yet behavioral extinction was rapid. Thus, the data did not support delayed priming.

## Discussion

This study tested whether naturally generated reward-omission dopamine pauses are necessary for behavioral extinction. Whereas prior studies largely examined the causal sufficiency of experimentally imposed dopamine-neuron excitation or inhibition^4,8–12^, we used a connectomic strategy to test the necessity of endogenous pauses. Gabazine^DART^ attenuated electrophysiologically defined pauses and photometrically measured reward-omission dopamine dips, yet accelerated rather than delayed behavioral extinction. Gabazine^DART^ also did not impair the decline of DA↑^cue*A*^, an internal measure of the learned *A*→reward association. Thus, intact natural reward-omission pauses were not necessary for either process.

These findings were not explained by generalized dopamine disruption or locomotor activation. Gabazine^DART^ spared tonic and burst firing, upward-going dopamine dynamics, and new cue–reward learning. Although gabazine^DART^ initially attenuated ITI and omission dips in parallel, the two signals subsequently diverged: ITI dip attenuation remained stable and predicted locomotor activation, whereas omission dips evolved and predicted behavioral extinction. Locomotion and extinction were also uncorrelated across the 39-mouse behavioral cohort. Together, these findings argue against generalized dopamine disruption, ITI dip attenuation, or locomotor activation as the explanation for accelerated behavioral extinction.

The temporal ordering further challenges the canonical view of naturally generated reward-omission pauses as necessary teaching signals for extinction^6–9^. The outcome-associated DA↑^reward*B*^ signal showed the expected teaching-signal sequence: it persisted while DA↑^cue*B*^ and cue-*B* prelicking emerged. By contrast, omission-related dynamics showed the reverse: the outcome-associated DA↓^omit^ signal dissipated before either DA↑^cue*A*^ declined or cue-*A* prelicking extinguished—opposite to a teaching signal that remains active until learning is established. The data also argue against a delayed-priming model in which omission dips initiate extinction but disappear before its behavioral expression: individuals with the most persistent DA↓^omit^ showed the longest post-dissipation delays, and several gabazine^DART^-treated mice showed little or no detectable DA↓^omit^ from the outset yet extinguished rapidly. Finally, gabazine^DART^ advanced both DA↓^omit^ dissipation and behavioral extinction while preserving their temporal relationship, consistent with advancing a shared transition rather than recruiting an independent route to behavioral extinction. A parsimonious interpretation is therefore that sustained DA↓^omit^ marks an extinction-resistant state, whereas its dissipation marks entry into an extinction-permissive state.

Notably, DA↓^omit^ tightly predicted behavioral extinction but was not detectably associated with DA↑^cue*A*^ decline. ANCCR (adjusted net contingency for causal relations), a retrospective causal-learning model, provides a useful framework for this dissociation^26^. ANCCR represents associations from outcomes back to preceding cues and updates cue←reward relationships only when rewards occur. Under this framework, training days 1-10 establish a strong *A*←reward retrospective association (“reward is always preceded by cue *A*”). After the rule flip on day 11, rewards would both gradually establish *B*←reward (“reward is now preceded by cue *B*”) and erase *A*←reward (“reward is no longer preceded by cue *A*”). However, a critical constraint of this framework is that *A*←reward cannot be updated on unrewarded trials. Consequently, this model predicts that unexpected omissions generate a competing *A*←omission association. Accordingly, ANCCR predicts that behavioral extinction in our task can arise through two independent pathways: *A*←omission learning and *A*←reward erasure, of which only *A*←omission learning should be directly modulated by omission-associated dopamine dips^24^. Under this framework, our data suggest that gabazine^DART^ accelerates DA↓^omit^ dissipation and *A*←omission learning, which appears to dominate behavioral extinction in this task. Conversely, DA↑^cue*A*^ decline may reflect *A*←reward erasure, which proceeds independently and on a generally slower timescale.

This interpretation preserves an important causal boundary. Gabazine^DART^ attenuated inhibitory input onto dopamine neurons and advanced a neural transition marked by DA↓^omit^ dissipation, but DA↓^omit^ need not itself be the sole causal variable: it could permit *A*←omission learning or report a deeper circuit transition that does so. Gabazine^DART^ may also shift the balance among parallel extinction mechanisms. For example, DA↓^omit^ dissipation could mark the point at which *A*←omission learning overtakes *A*←reward erasure. The causal conclusion should therefore remain at the level directly supported by the experiment: attenuating inhibitory input onto dopamine neurons advances a neural transition marked by DA↓^omit^ dissipation and accelerates behavioral extinction.

Prior studies bearing on necessity provide relevant but non-equivalent context. Calibrated excitation of dopamine neurons at reward omission did not accelerate behavioral extinction^4^, underscoring that insertion of exogenously timed excitation is not the same as attenuating endogenous inhibitory inputs. Chronic knockout of a GABA_A_ receptor subunit in dopamine neurons likewise did not accelerate extinction^17^, highlighting the importance of acute perturbations to minimize circuit compensation. Finally, lesions of the habenular complex reduced dopamine-neuron inhibition following reward omission while sparing inhibition to air-puff events; these lesions altered learning in a manner consistent with reduced negative relative to positive reward-prediction-error signaling, seemingly at odds with our findings^3^. However, their task used probabilistic conditioning rather than extinction following a discrete switch to reward omission, and the lesion altered multiple components of VTA reward-prediction-error circuitry and could influence behavior through habenular projections to non-dopaminergic circuits.

Our results further complement prior sufficiency studies showing that experimentally imposed dopamine-neuron inhibition can counteract the reinforcing effects of reward delivery^9–11^. Sufficiency and necessity address distinct causal questions: an imposed pause may be sufficient to reduce behavioral responding even when intact naturally generated omission pauses are not required for extinction. The effects of imposed inhibition may also depend on circuit origin, timing, and behavioral context. A recent preprint illustrates this context dependence: optogenetically augmenting dorsomedial-striatal dopamine dips after unrewarded actions increased subsequent reward seeking despite punishment^27^. The preprint tested the sufficiency of dopamine dips in punishment-resistant reward seeking, whereas our work tested the necessity of natural omission pauses during Pavlovian extinction. Nevertheless, the results are directionally aligned: augmenting dopamine dips promoted persistence, whereas attenuating pause-generating inhibitory input reduced persistence. Collectively, these studies support a broader conclusion: dopamine-neuron pauses are not a unitary signal with a fixed function across circuit loci, tasks, and behavioral states.

Several limitations remain. First, gabazine^DART^ broadly attenuates GABA_A_-mediated inhibition onto dopamine neurons rather than selectively targeting the inputs recruited by reward omission. The double dissociation between DA↓^ITI^ and DA↓^omit^ dynamics narrows this concern, but tools with greater specificity will be needed to identify the relevant presynaptic sources. Second, population photometry cannot identify the dopamine-neuron subtype or projection-defined channel contributing to the extinction-linked signal. Third, the temporal and correlational dissociations are consistent with distinct A←omission learning and A←reward erasure, but do not uniquely establish this decomposition; we therefore treat it as an interpretive framework rather than a measured separation of memory traces. Finally, the task combined extinction of cue A with conditioning of a newly rewarded cue B. This design maintained reward availability and provided a within-animal control for new learning, but confines the present conclusions to extinction during reward reassignment.

Behavioral persistence is adaptive while established associations remain reliable but can become maladaptive as contingencies change. By showing that targeted attenuation of inhibitory input onto dopamine neurons accelerates extinction without disrupting upward-going dopamine dynamics or new learning, these findings show that pause-generating inhibitory input sustains behavioral persistence and governs when behavior becomes flexible. More broadly, the results illustrate how connectomic perturbations can reveal the functions of endogenous neural motifs.

---

**fig. S1.**
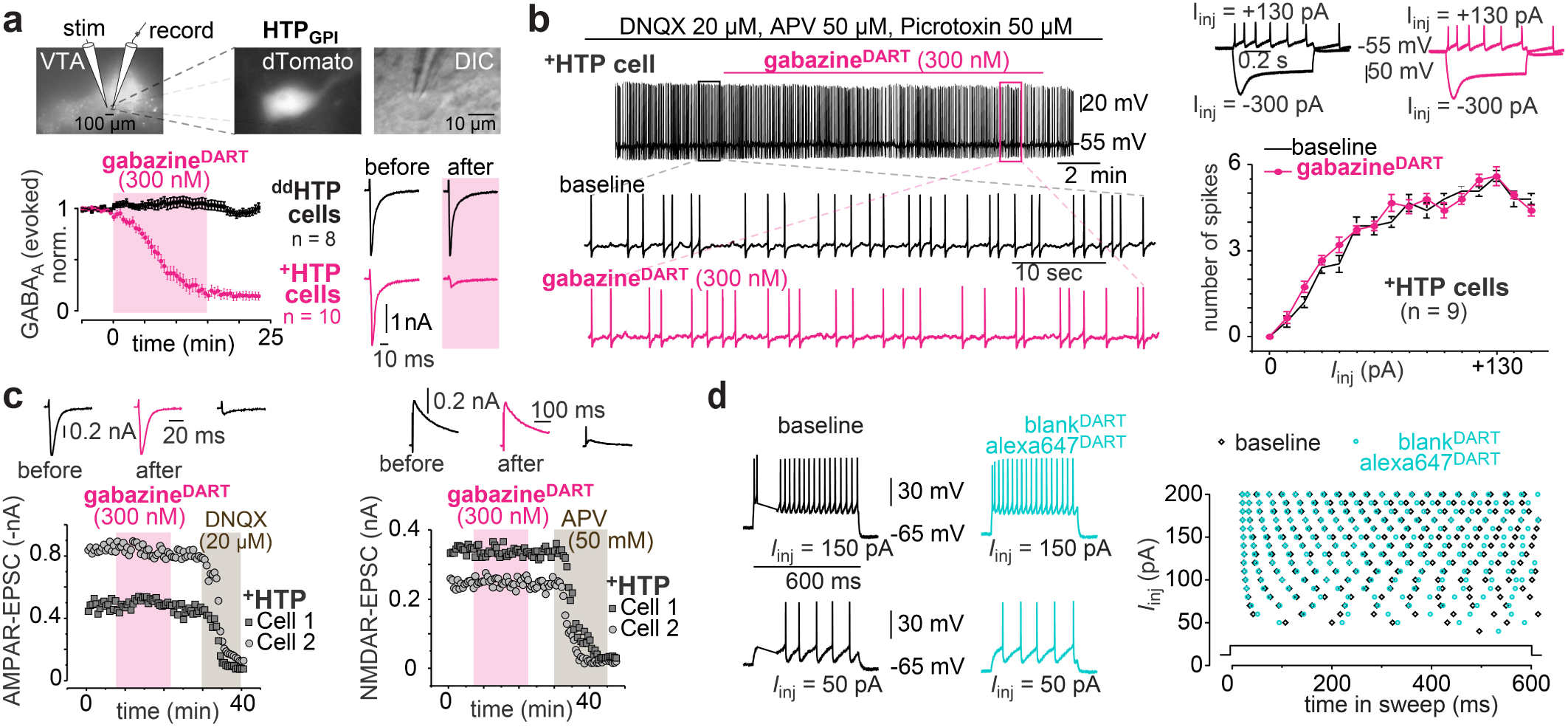
Gabazine^DART^ enables acute, cell-type-specific antagonism of the GABAAR. (a) Gabazine^DART^ blocks evoked GABA_A_R currents in ^+^HTP-expressing VTA^DA^ neurons with no impact on neurons expressing a control ^dd^HTP construct. Top: evoked-IPSC configuration. Bottom: 300 nM gabazine^DART^ has no impact on ^dd^HTP neurons, while blocking IPSCs on ^+^HTP cells in under 15 min (86 ± 4% block). Data are mean ±SEM, cells normalized to baseline (^+^HTP: n=10 cells; ^dd^HTP: n=8 cells). Example traces to right. (b) Gabazine^DART^ does not impact VTA_DA_ action potentials. Left: current clamp of a VTA_DA_ neuron in the presence of picrotoxin (GABA_A_ blocker), DNQX (AMPA blocker), and APV (NMDA blocker). Right: quantification of action potential firing as a function of injected current; performed before (black) vs after (cyan) gabazine^DART^ was tethered on each cell. Representative traces shown above. Error bars are mean ±SEM over cells (*n* = 9). (c) Gabazine^DART^ does not impact AMPA or NMDA receptors. Left: 300 nM gabazine^DART^ has no effect on ^+^HTP neuron AMPAR-EPSCs, which are subsequently blocked by 20 µM DNQX. Example traces above. Right: 300 nM gabazine^DART^ has no effect on ^+^HTP neuron NMDAR-EPSCs, which are subsequently blocked by 50 µM APV. Example traces above. (d) Alexa647^DART^ is pharmacologically inert. Current clamp studies in VTA_DA_ ^+^HTP neurons with 10:1 blank^DART^ + alexa647^DART^. No significant change was observed before vs after alexa647^DART^ was tethered. Representative traces shown left.

**fig. S2.**
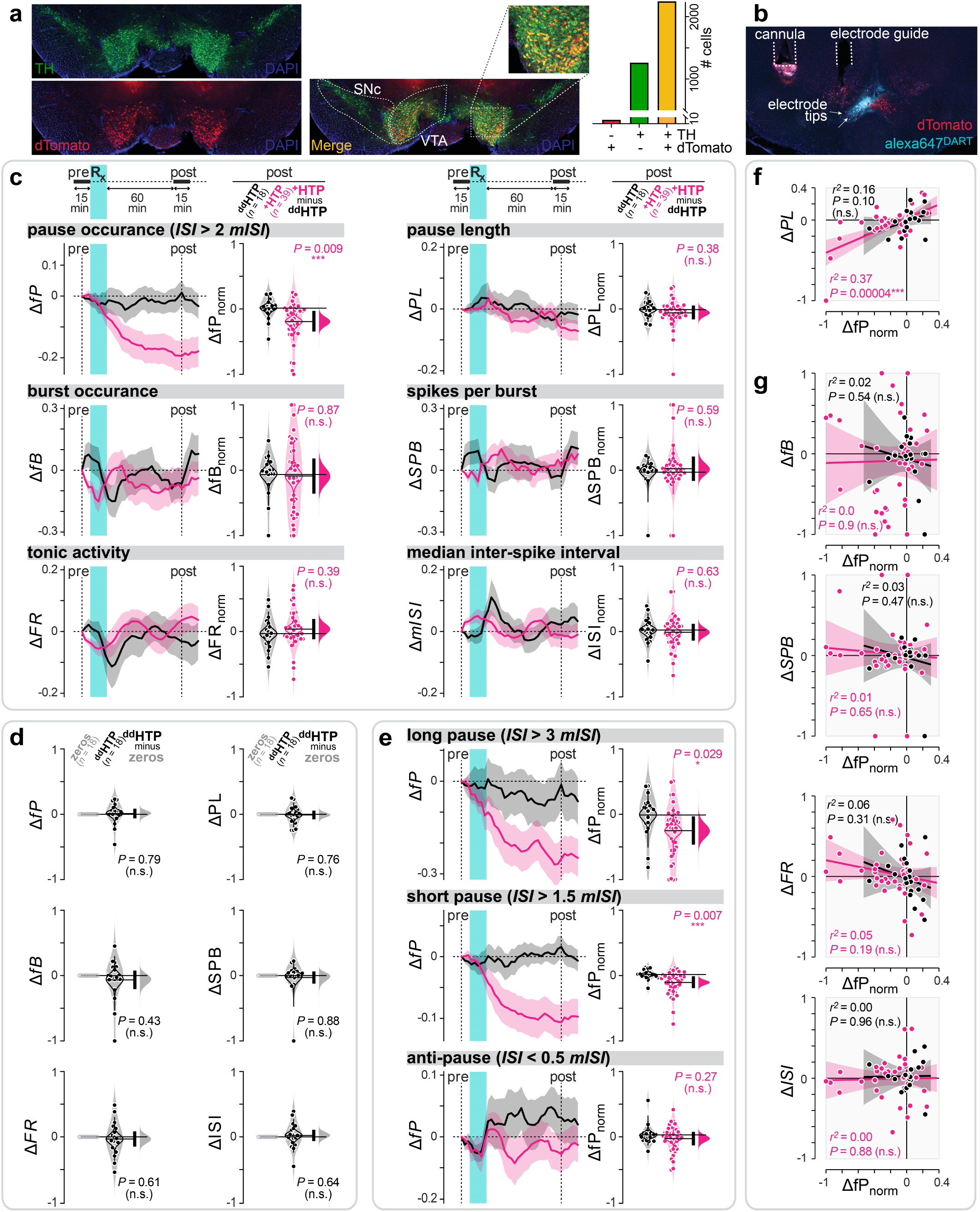
Gabazine^DART^ only blocks dopamine pauses. (a) Representative histology and quantitative cell counting. Dopamine neurons (TH, tyrosine-hydroxylase, green). HTP expression (dTomato, red). Cell counting was performed from a representative brain containing 2,244 double-labeled (dTomato^+^/TH^+^) cells. This represents 99.7% of all virus-positive cells (2,250 dTomato^+^), and 64% of all dopaminergic cells (3,507 TH^+^). (b) Electrode histology. Post-electrophysiology histology. HTP expression (dTomato, red); ligand capture (cyan); and electrode tips (arrows). (c) Gabazine^DART^ selectively reduces the occurrence of VTA^DA^ pauses without impacting burst or tonic firing. Top row, left: pause occurrence, *fP* (fraction of *ISI* >2× the median). The 15-min baseline (*fP*_pre_) is compared to a sliding 15-min window (*fP*_post_) using Δ*fP* = (*fP*_post_ *-fP*_pre_) / (*fP*_post_ + *fP*_pre_). Shading indicates mean ± SEM across cells (^dd^HTP: *n* = 18 cells, 3 mice; ^+^HTP: *n* = 39 cells, 5 mice). Right: at 1-hr post-gabazine^DART^, Δ*fP* differed significantly between ^+^HTP and ^dd^HTP cells. Individual cells (dots), kernel density estimate (shaded violin), mean bootstrap (outlined violin), difference bootstrap (pink distribution), and 95% CI of the two-sided permutation test (vertical black bar; *P* = 0.009). Other metrics: *fB* (fraction of spikes fired in bursts), *FR* (firing rate), *PL* (pause length), *SPB* (spikes per burst), and *mISI* (median interspike interval) revealed no significant ^+^HTP vs ^dd^HTP differences. (d) Gabazine^DART^ has no impact in ^dd^HTP mice. Steady-state Δ (comparing ^dd^HTP cells to zero, 1-hr post-gabazine^DART^) with individual cells (dots), kernel density estimate (shaded violin), mean bootstrap (outlined violin), difference bootstrap (grey distribution), and 95% CI of the two-sided permutation test (vertical black bar). (e) Robustness analysis of the primary pause metric, *fP*. The top two panels examine two alternate definitions of a pause (fraction *ISI* > 3 × the median; fraction *ISI* > 1.5 × the median); in both cases, gabazine^DART^ produced a significant reduction in pauses in ^+^HTP vs ^dd^HTP mice, congruent with the standard *fP* (fraction *ISI* > 2 × the median). The bottom panel examines an ‘anti-pause’ (fraction *ISI* < 0.5 × the median), where the lack of difference in ^+^HTP vs ^dd^HTP cells suggests that changes cannot be explained by a symmetrical change in *ISI* variance. (f) Pauses vs pause length. Correlation between *PL* (pause length) and *fP* from each ^+^HTP cell (circles, *n*=39), with regression ±95% CI (line and shading). Pearson’s *r*^2^ = 0.37, *P* = 0.00004. (g) Pauses vs other features. Correlation between all other metrics and *fP*; format as above.

**fig. S3.**
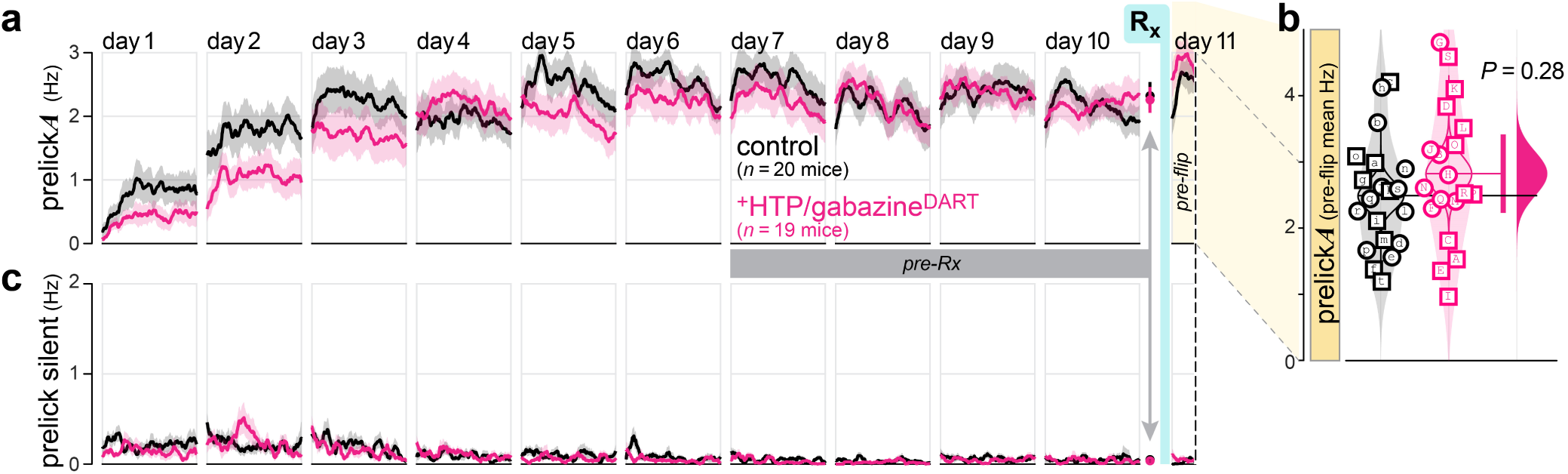
Gabazine^DART^ does not alter cue-*A* prelicking before the rule flip. **(a)** Cue-*A* prelicking was acquired during *A*→reward training and remained stable after infusion, before the rule flip. Raw cue-*A* prelicking rates (in Hz) are shown across training days 1–10 and the 15-min pre-flip period on day 11. The cyan bar marks infusion of gabazine^DART^ and alexa647^DART^, which occurred 2 h before the behavioral session on day 11. Lines and shading indicate mean ± s.e.m. Pink, ^+^HTP/gabazine^DART^ (n = 19 mice); black, controls (*n* = 20 mice). **(b)** Mean cue-*A* prelicking during the pre-flip period did not differ detectably between ^+^HTP and ^dd^HTP mice (*P* = 0.28, two-sided permutation test). Shown are individual mice (male, square; female, circle), kernel density estimates (shaded violins), bootstrap distributions of the group means (outlined violins), and group means (horizontal lines). At right, hypothesis testing is shown as the 95% CI from a two-sided permutation test (vertical pink bar), and effect size as the bootstrap distribution of the mean difference (pink half-violin). Lettered symbols identify the same mice throughout the manuscript. **(c)** Prelicking during silent trials diminished towards zero over training. Raw silent-trial prelicking rates are shown across the same days and time periods as in **a**. Lines and shading indicate mean ± s.e.m.

**fig. S4.**
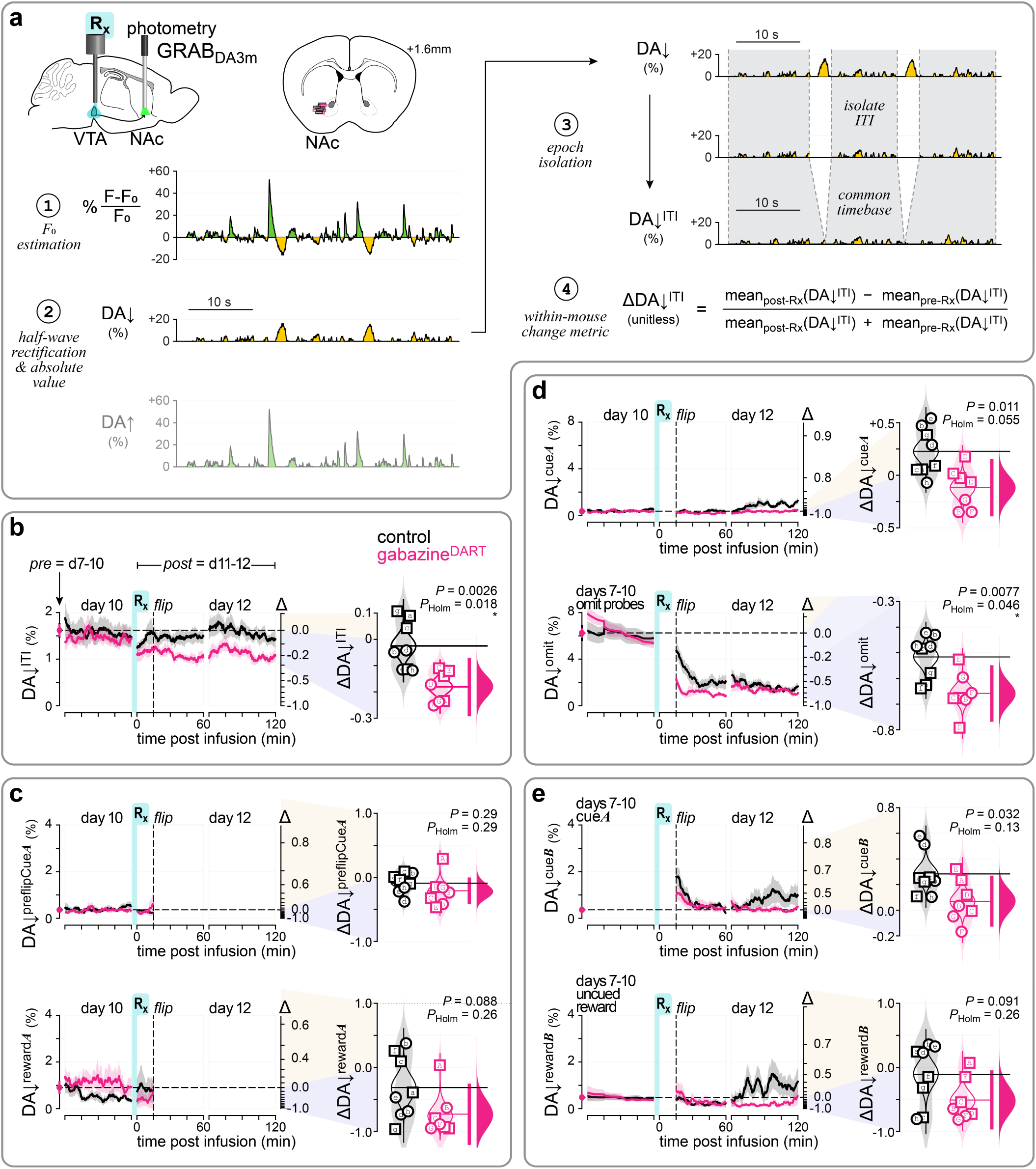
Gabazine^DART^ attenuates downward-going dopamine signals during ITIs and reward omission. **(a)** Rectified epoch-analysis workflow for nucleus accumbens GRAB_DA3m_ photometry. Gabazine^DART^ was infused into the VTA while dopamine fluorescence was recorded in the lateral nucleus accumbens. **Step 1**: baseline fluorescence, *F*_0_, was estimated using an iterative moving-median procedure (**methods**) and used to calculate percentage fluorescence change, (*F* − *F*_0_) / *F*_0_. **Step 2**: the continuous fluorescence signal was separated by half-wave rectification into downward-going and upward-going components. Samples of the opposite polarity were set to zero but retained in the continuous trace, ensuring that each component captured both the amplitude and prevalence of deflections in the analyzed direction. The downward-going component was sign-inverted so that DA↓ and DA↑ were both non-negative. **Step 3**: each component was isolated within predefined behavioral epochs and placed on a common time base. The example shows DA↓ during intertrial intervals (DA↓^ITI^). **Step 4**: within-mouse change was calculated as Δ = (post − pre) / (post + pre), using the mean signal within the corresponding pre-and post-infusion periods. **(b)** Gabazine^DART^ attenuated DA↓^ITI^. **Left,** baseline-aligned signal amplitude in native percentage fluorescence-change units. Lines and shading indicate mean ± s.e.m. for *n* = 7 ^+^HTP/gabazine^DART^ mice (pink) and *n* = 8 control mice (black). For display only, data from each treatment group were multiplied by a single constant so that the day 7–10 pre-infusion time average for that group equaled the geometric mean of the two group baselines (far-left symbol and dashed horizontal line). This group-level rescaling preserved relative within-group variability without normalizing individual mice. Raw values are provided in supplementary data. The right axis shows Δ = (post − pre) / (post + pre), comparing the post-infusion waveform to the pre-infusion mean from days 7–10. More negative Δ values indicate greater post-infusion attenuation of downward-going signals. **Right,** ΔDA↓^ITI^ differed significantly between ^+^HTP/gabazine^DART^ and control mice (*P*_unadj_ = 0.0026, *P*_Holm_ = 0.018). Shown are individual mice (male, square; female, circle), kernel density estimates (shaded violins), bootstrap distributions of the group means (outlined violins), and group means (horizontal lines). At right, hypothesis testing is shown as the 95% CI from a two-sided permutation test (vertical pink bar), and effect size as the bootstrap distribution of the mean difference (pink half-violin). Lettered symbols identify the same mice throughout the manuscript. Holm correction was applied across the seven predefined behavioral epochs. **(c)** Gabazine^DART^ did not detectably alter downward-going signals during the pre-flip cue-*A* or reward-*A* epochs. **Top,** DA↓^preflipCue*A*^ before the rule flip, using DA↓^cue*A*^ from training days 7–10 as the pre-infusion reference (*P*_unadj_ = 0.29, *P*_Holm_ = 0.29). **Bottom,** DA↓^reward*A*^ before the rule flip, using DA↓^reward*A*^ from training days 7–10 as the pre-infusion reference (*P*_unadj_ = 0.088, *P*_Holm_ = 0.26). Format as in **b**. **(d)** Gabazine^DART^ attenuated DA↓^omit^, whereas attenuation of DA↓^cue*A*^ did not survive family-wise correction. **Top,** DA↓^cue*A*^ after the rule flip, using DA↓^cue*A*^ from training days 7–10 as the pre-infusion reference (*P*_unadj_ = 0.011, *P*_Holm_ = 0.055). **Bottom,** DA↓^omit^ after the rule flip, using probe omissions on training days 7–10 as the pre-infusion reference (*P*_unadj_ = 0.0077, *P*_Holm_ = 0.046). Time courses and full-period change metrics are displayed as in **b**. **(e)** Gabazine^DART^ did not detectably alter downward-going signals during cue-*B* or reward-*B* epochs after family-wise correction. **Top,** DA↓^cue*B*^ after the rule flip, using DA↓^cue*A*^ from training days 7–10 as the pre-infusion reference (*P*_unadj_ = 0.032, *P*_Holm_ = 0.13). **Bottom,** DA↓^rewardB^ after the rule flip, using responses to uncued rewards on training days 7–10 as the pre-infusion reference (*P*_unadj_ = 0.091, *P*_Holm_ = 0.26). Time courses and full-period change metrics are displayed as in **b**.

**fig. S5.**
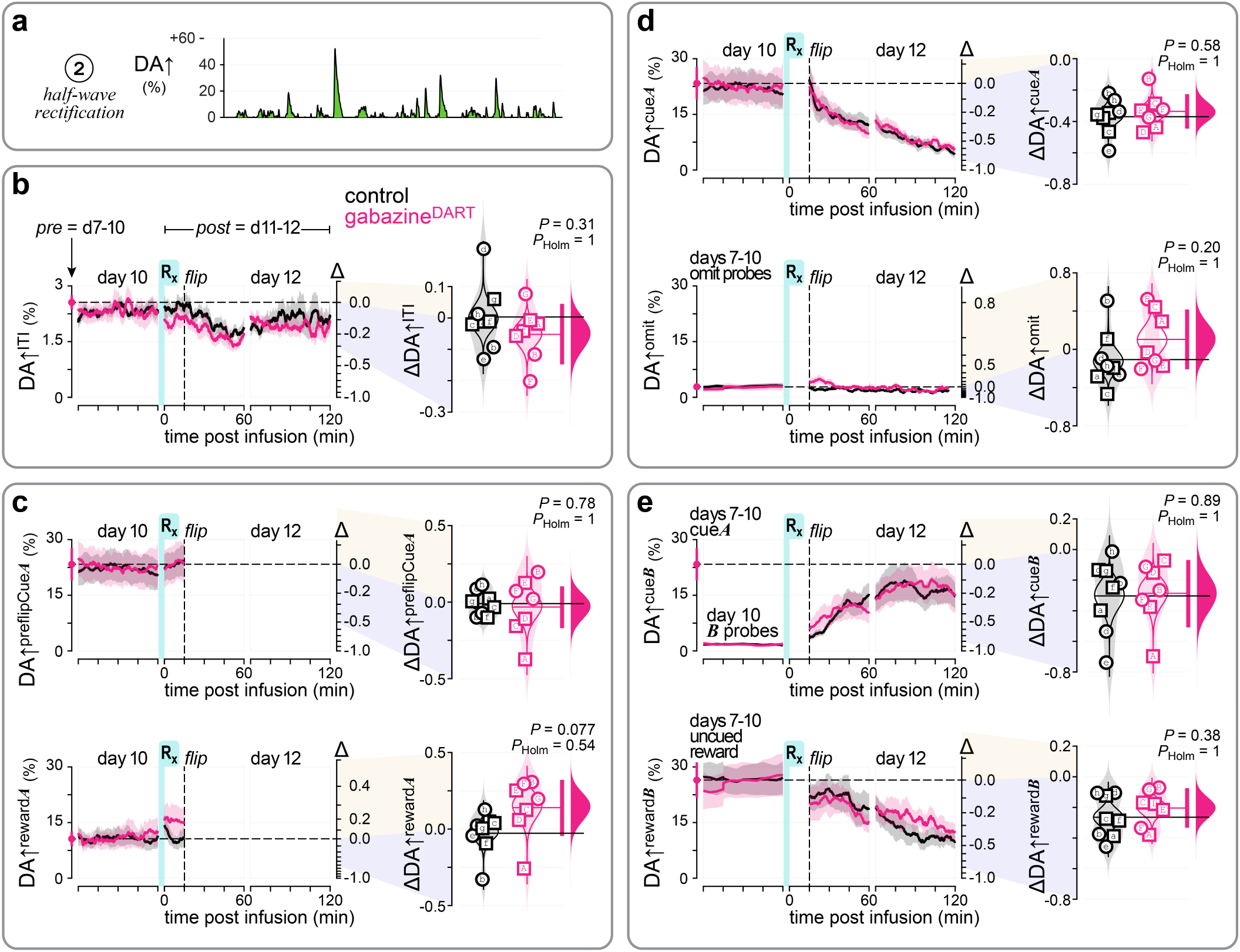
Gabazine^DART^ does not detectably alter upward-going dopamine signals across behavioral epochs. **(a)** Half-wave rectification isolated upward-going dopamine signals, as described in **fig. S4a**. **(b)** Gabazine^DART^ did not detectably alter DA↑^ITI^. **Left,** baseline-aligned signal amplitude in native percentage fluorescence-change units. Lines and shading indicate mean ± s.e.m. for *n* = 7 ^+^HTP/gabazine^DART^ mice (pink) and *n* = 8 control mice (black). The right axis shows Δ = (post − pre) / (post + pre), comparing the post-infusion waveform with the pre-infusion mean from days 7–10. **Right,** full-period ΔDA↑^ITI^ did not differ detectably between ^+^HTP/gabazine^DART^ and control mice (*P*_unadj_ = 0.31, *P*_Holm_ = 1). Shown are individual mice (male, square; female, circle), kernel density estimates (shaded violins), bootstrap distributions of the group means (outlined violins), and group means (horizontal lines). At right, hypothesis testing is shown as the 95% CI from a two-sided permutation test (vertical pink bar), and effect size as the bootstrap distribution of the mean difference (pink half-violin). Holm correction was applied across the seven predefined behavioral epochs. **(c)** Gabazine^DART^ did not alter upward-going signals during the pre-flip cue-*A* or reward-*A* epochs. **Top,** DA↑^preflipCue*A*^ before the rule flip, using DA↑^cue*A*^ from days 7–10 as the pre-infusion reference (*P*_unadj_ = 0.78, *P*_Holm_ = 1). Bottom, DA↑^reward*A*^ before the rule flip, using DA↑^reward*A*^ from days 7–10 as the pre-infusion reference (*P*_unadj_ = 0.077, *P*_Holm_ = 0.54). Format as in **b**. **(d)** Gabazine^DART^ did not detectably alter DA↑^cue*A*^ or DA↑^omit^ after the rule flip. **Top,** DA↑^cue*A*^ after the rule flip, using DA↑^cue*A*^ from days 7–10 as the pre-infusion reference (*P*_unadj_ = 0.58, *P*_Holm_ = 1). **Bottom,** DA↑^omit^ after the rule flip, using probe omissions on days 7–10 as the pre-infusion reference (*P*_unadj_ = 0.20, *P*_Holm_ = 1). Format as in **b**. **(e)** Gabazine^DART^ did not detectably alter upward-going signals during cue-*B* or reward-*B* epochs. **Top,** DA↑^cue*B*^ after the rule flip, using DA↑^cue*A*^ from days 7–10 as the pre-infusion reference; responses to five unrewarded cue-B probes on day 10 are shown immediately before infusion (*P*_unadj_ = 0.89, *P*_Holm_ = 1). **Bottom,** DA↑^reward*B*^ after the rule flip, using responses to uncued rewards on days 7–10 as the pre-infusion reference (*P*_unadj_ = 0.38, *P*_Holm_ = 1). Format as in **b**.

**fig. S6.**
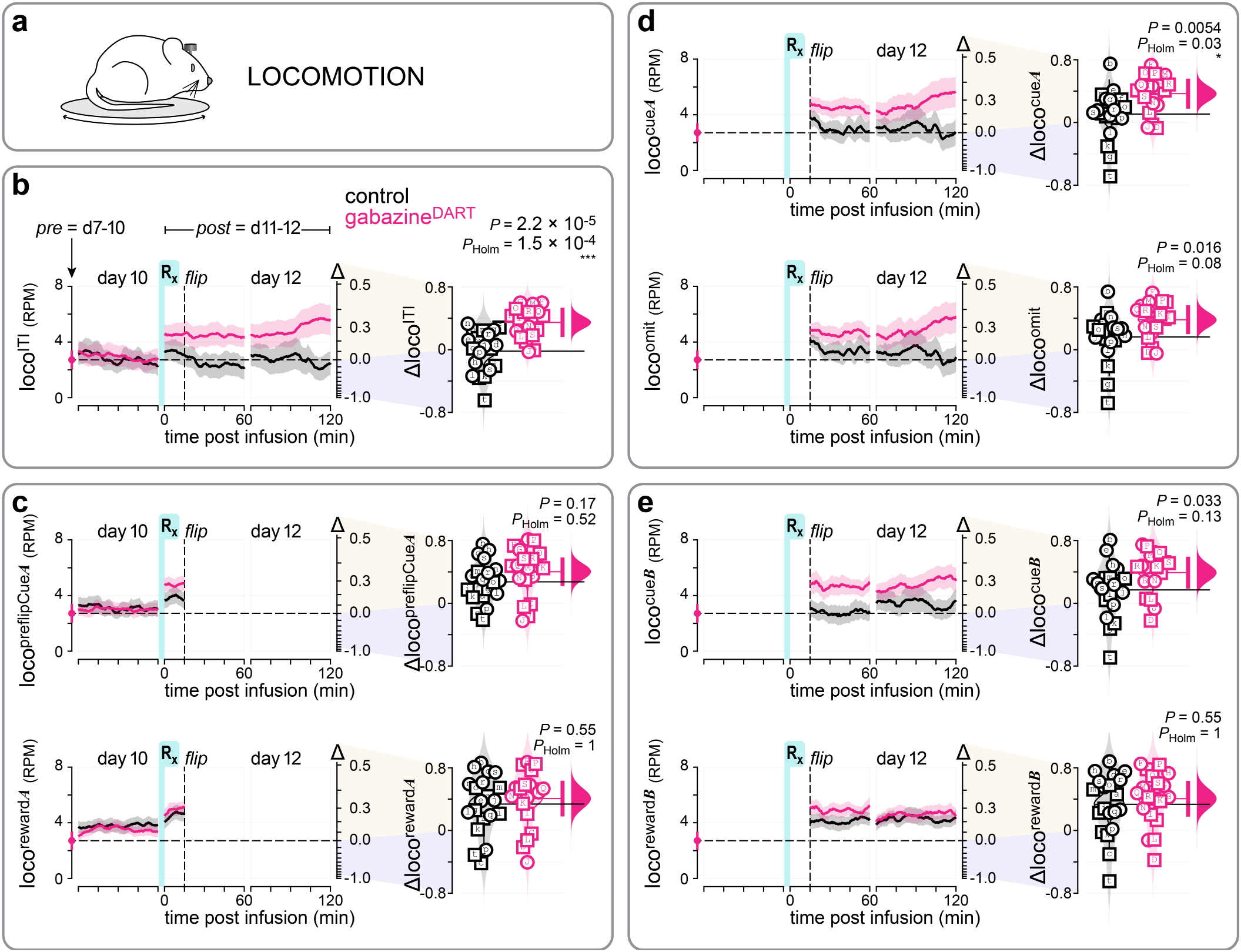
Gabazine^DART^ increases locomotion during ITIs and post-flip cue-A epochs. **(a)** Locomotion was measured from running-wheel rotations during head-fixed behavior. **(b)** Gabazine^DART^ increased locomotion during intertrial intervals. **Left,** baseline-aligned ITI locomotion (loco^ITI^) in wheel rotations per min across days 10–12. Lines and shading indicate mean ± s.e.m. for *n* = 19 ^+^HTP/gabazine^DART^ mice (pink) and *n* = 20 control mice (black). For display only, data from each treatment group were multiplied by a single constant so that the day 7–10 pre-infusion loco^ITI^ average for that group equaled the geometric mean of the two group baselines (far-left symbol and dashed horizontal line). This group-level rescaling preserved relative within-group variability without normalizing individual mice. Raw values are provided in supplementary data. For all locomotion epochs in **b–e**, the common pre-infusion reference was loco^ITI^ averaged across days 7–10; this reference was used because not all mice received probe omissions or uncued rewards. **Right,** Δloco^ITI^ was greater in ^+^HTP/gabazine^DART^ than control mice (*P*_unadj_ = 2.2 × 10⁻⁵, *P*_Holm_ = 1.5 × 10⁻⁴). Shown are individual mice (male, square; female, circle), kernel density estimates (shaded violins), bootstrap distributions of the group means (outlined violins), and group means (horizontal lines). Hypothesis testing is shown as the 95% CI from a two-sided permutation test (vertical pink bar), and effect size as the bootstrap distribution of the mean difference (pink half-violin). Holm correction was applied across the seven predefined behavioral epochs. **(c)** Gabazine^DART^ did not alter locomotion during the pre-flip cue-*A* or reward-*A* epochs. **Top,** locomotion during pre-flip cue-*A* (*P*_unadj_ = 0.17, *P*_Holm_ = 0.52). **Bottom,** locomotion during pre-flip reward (*P*_unadj_ = 0.55, *P*_Holm_ = 1). Format as in **b**. **(d)** Gabazine^DART^ increased locomotion during post-flip cue-*A* presentation, whereas the increase during reward omission did not survive family-wise correction. **Top,** locomotion during cue-*A* presentation after the rule flip (*P*_unadj_ = 0.0054, *P*_Holm_ = 0.034). **Bottom,** locomotion during post-flip reward omission (*P*_unadj_ = 0.016, *P*_Holm_ = 0.08). Format as in **b**. **(e)** Gabazine^DART^ did not detectably alter locomotion during cue-*B* or reward-*B* epochs after family-wise correction. **Top,** locomotion during post-flip cue-*B* presentation (*P*_unadj_ = 0.033, *P*_Holm_ = 0.13). **Bottom,** locomotion during post-flip reward-*B* delivery (*P*_unadj_ = 0.55, *P*_Holm_ = 1). Format as in **b**.

**fig. S7.**
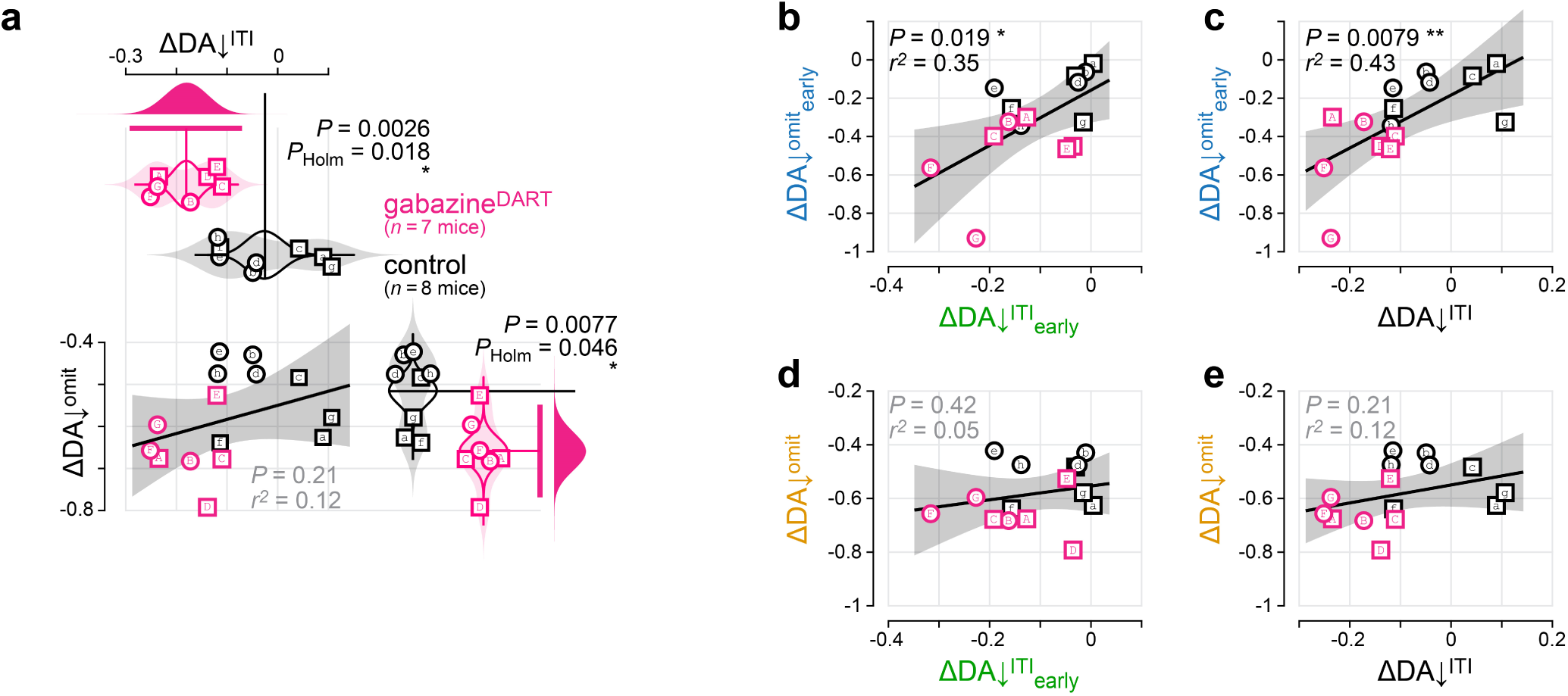
ΔDA↓^omit^early—but not full-period ΔDA↓^omit^—covaries with ΔDA↓^ITI^. **(a)** Gabazine^DART^ attenuated both DA↓^ITI^ and DA↓^omit^ across the full post-infusion period, but the two changes did not detectably covary across mice. **Top,** full-period ΔDA↓^ITI^ differed between ^+^HTP/gabazine^DART^ and control mice (*P*_unadj_ = 0.0026, *P*_Holm_ = 0.018). **Bottom left,** relationship between full-period ΔDA↓^ITI^ and ΔDA↓^omit^ (Pearson’s *P* = 0.21, *r*² = 0.12). **Bottom right,** full-period ΔDA↓^omit^ differed between groups (*P*_unadj_ = 0.0077, *P*_Holm_ = 0.046). Full-period values compare the fixed pre-infusion mean with the mean post-infusion signal from 0–120 min. Group comparisons are displayed as in **fig. S4b**. Solid lines and shading indicate linear fits and 95% confidence intervals. Pink, ^+^HTP/gabazineDART (*n* = 7 mice); black, control (*n* = 8 mice). Lettered symbols identify the same mice throughout the manuscript. **(b)** ÄDA↓^omit^_early_ covaried with ΔDA↓^ITI^_early_ (Pearson’s *P* = 0.019, *r*² = 0.35). ΔDA↓^omit^_early_ compares the pre-infusion mean with the mean from the first four post-flip omissions. ΔDA↓^ITI^_early_ compares the pre-infusion mean with the mean from 0–15 min post-infusion. Solid lines and shading indicate linear fits and 95% confidence intervals. **(c)** ÄDA↓^omit^_early_ also covaried with full-period ΔDA↓^ITI^ (Pearson’s *P* = 0.0079, *r*² = 0.43). Format as in **b**. **(d)** Full-period ΔDA↓^omit^ did not detectably covary with ΔDA↓^ITI^_early_ (Pearson’s *P* = 0.42, *r*² = 0.05). Format as in **b**. **(e)** Full-period ΔDA↓^omit^ did not detectably covary with full-period ΔDA↓^ITI^ (Pearson’s *P* = 0.21, *r*² = 0.12). Format as in **b**.

**fig. S8.**
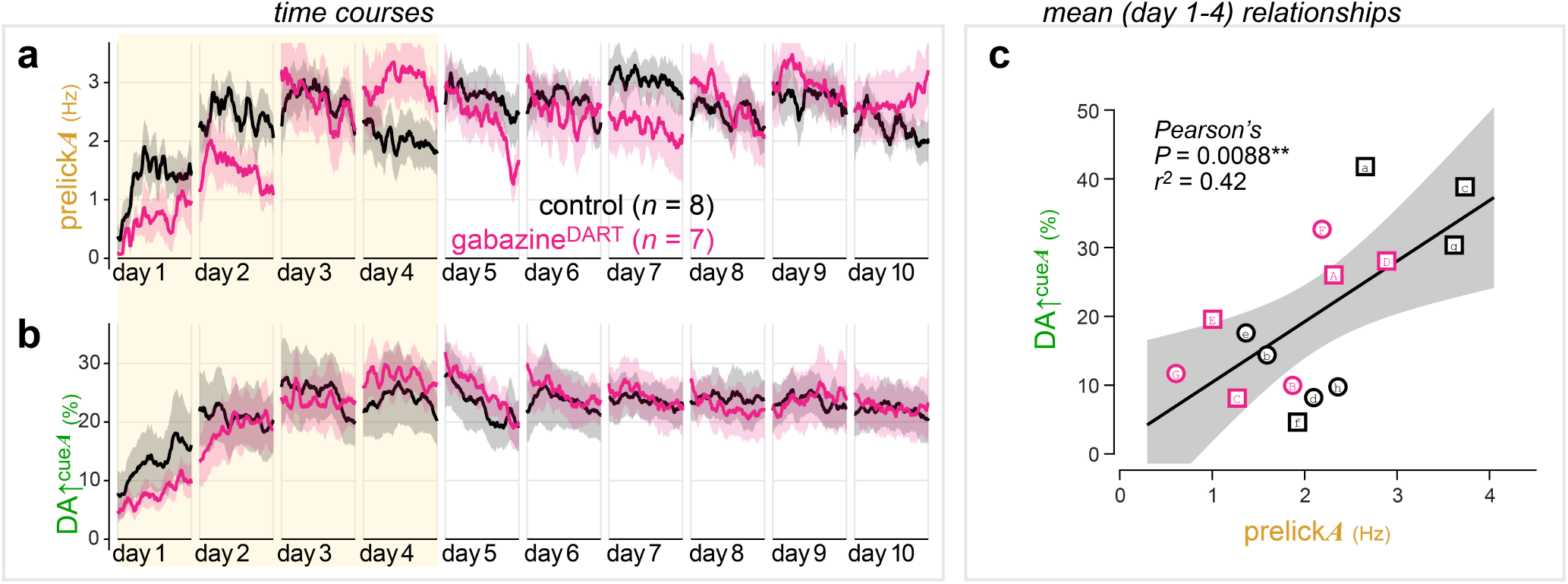
Cue-A prelicking and DA↑^cue*A*^ emerge together during *A*→reward training. **(a)** Cue-*A* prelicking developed during the first four days of *A*→reward training and remained robust through day 10. Raw cue-*A* prelicking rates are shown across training days 1–10. Lines and shading indicate mean ± s.e.m. for mice subsequently assigned to control (black; *n* = 8 mice) or ^+^HTP/gabazine^DART^ (pink; *n* = 7 mice) groups. All data were collected before treatment. Yellow shading marks training days 1–4, used for the analysis in **c**. **(b)** The internal DA↑^cue*A*^ signal also developed during the first four days of *A*→reward training. Upward-going nucleus accumbens dopamine signals during cue-*A* presentation are shown across training days 1–10. Lines and shading indicate mean ± s.e.m.; groups and shading are as in **a**. **(c)** DA^↑cue*A*^ and cue-*A* prelicking were positively associated during early training. Each point shows one mouse’s mean DA↑^cue*A*^ and cue-*A* prelicking across training days 1–4 (Pearson’s *P* = 0.0088, *r*² = 0.42; *n* = 15 mice). Symbols indicate individual mice (male, square; female, circle); pink and black denote subsequent ^+^HTP/gabazine^DART^ and control assignments, respectively. The line and shading indicate the linear fit and 95% confidence interval.

**fig. S9.**
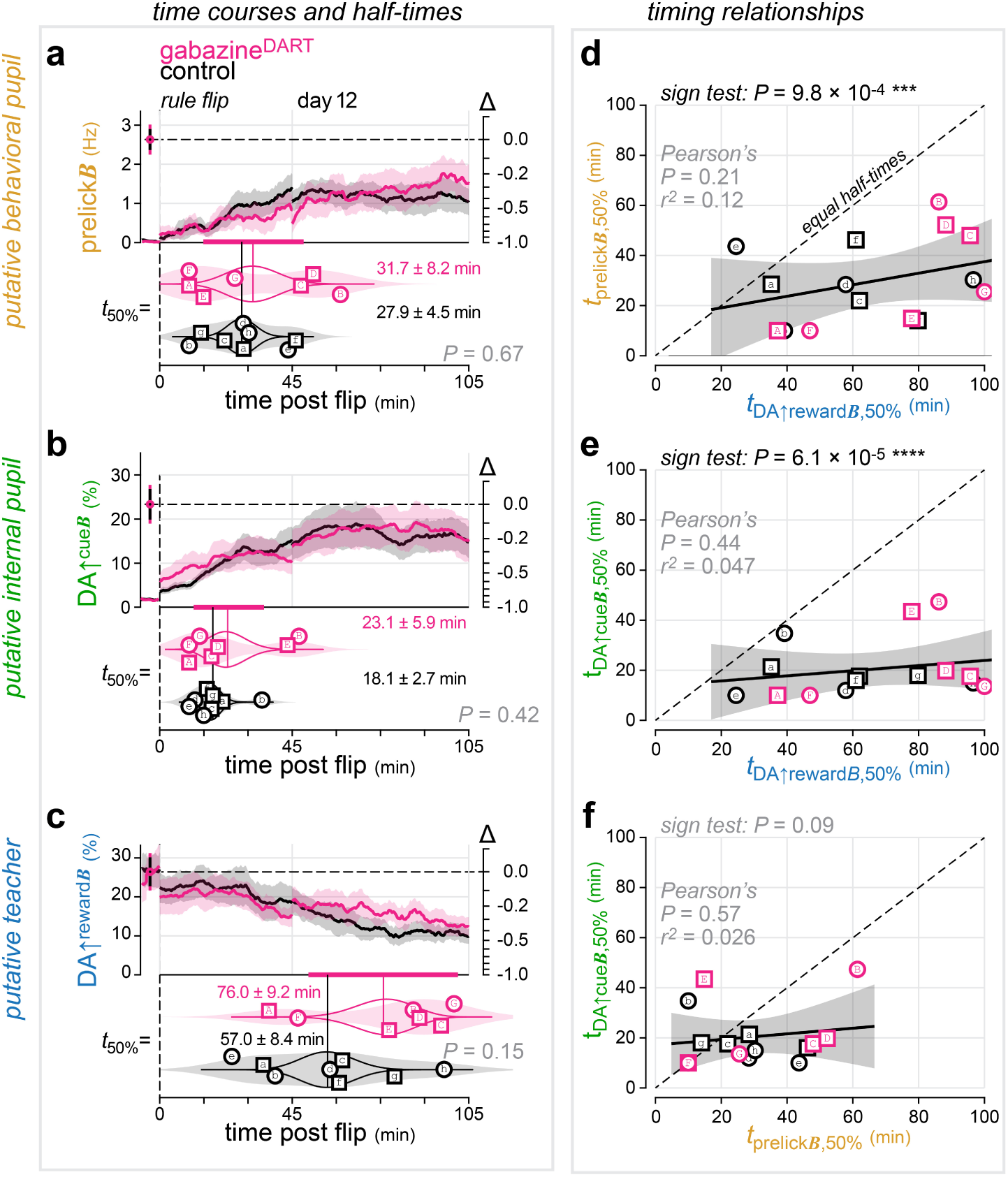
Gabazine^DART^ spares behavioral conditioning and cue-evoked dopamine emergence, and both occur before dissipation of the putative DA↑^rewardB^ teacher. **(a)** Gabazine^DART^ did not detectably alter behavioral conditioning to cue *B* within the photometry cohort. **Top,** cue-*B* prelicking across the post-flip period on days 11 and 12. The far-left symbols and dashed horizontal line show pre-flip cue-*A* prelicking, used as the reference for the normalized change shown on the right axis. Lines and shading indicate mean ± s.e.m. for *n* = 7 ^+^HTP/gabazine^DART^ mice (pink) and *n* = 8 control mice (black). **Bottom,** half-conditioning times (*t*_50%_) estimated from logistic fits for individual mice. Shown are individual mice (male, square; female, circle), kernel density estimates (shaded violins), bootstrap distributions of the group means (outlined violins), and group means (vertical lines). Hypothesis testing is shown as the 95% CI from a two-sided permutation test (horizontal pink bar). Half-conditioning times did not differ detectably between ^+^HTP/gabazine^DART^ mice (31.7 ± 8.2 min) and controls (27.9 ± 4.5 min; *P* = 0.67). **(b)** Gabazine^DART^ did not detectably alter emergence of DA↑^cue*B*^. Top, DA↑^cue*B*^ across the post-flip period. The far-left symbols and dashed horizontal line show DA↑^cue*A*^ averaged across training days 7–10, used as the pre-infusion reference for the normalized change shown on the right axis. Responses to five unrewarded cue-*B* probes on day 10 are shown immediately before infusion. Photometry time courses are displayed as described in Fig. 3a. **Bottom,** half-conditioning times estimated from logistic fits, displayed as in **a**. Half-times did not differ detectably between ^+^HTP/gabazine^DART^ mice (23.1 ± 5.9 min) and controls (18.1 ± 2.7 min; *P* = 0.42, two-sided permutation test). **(c)** Gabazine^DART^ did not detectably alter persistence of DA↑^reward*B*^. Top, DA↑^reward*B*^ across the post-flip period. The far-left symbols and dashed horizontal line show responses to uncued rewards averaged across training days 7–10, used as the pre-infusion reference for the normalized change shown on the right axis. **Bottom,** half-dissipation times estimated from logistic fits, displayed as in **a**. Half-times did not differ detectably between ^+^HTP/gabazine^DART^ mice (76.0 ± 9.2 min) and controls (57.0 ± 8.4 min; *P* = 0.15, two-sided permutation test). **(d)** Cue-*B* prelicking occurred before DA↑^reward*B*^ dissipation in 14 of 15 mice. The half-conditioning time of cue-*B* prelicking is plotted against the half-dissipation time of DA↑^reward*B*^. Pink and black symbols denote ^+^HTP/gabazine^DART^ and control mice, respectively; lettered symbols identify the same mice throughout the manuscript. Solid lines and shading indicate linear fits and 95% confidence intervals; dashed diagonals indicate equal half-times. Fourteen mice fall below the dashed line (two-sided sign test, *P* = 9.8 × 10⁻⁴). The two half-times were not detectably associated across mice (Pearson’s *P* = 0.21, *r*² = 0.12). **(e)** DA↑^cue*B*^ emerged before DA↑^reward*B*^ dissipation in every mouse. The half-conditioning time of DA↑^cue*B*^ is plotted against the half-dissipation time of DA↑^reward*B*^. All 15 mice fall below the dashed line of equal half-times (two-sided sign test, *P* = 6.1 × 10⁻⁵). The half-times were not detectably associated across mice (Pearson’s *P* = 0.44, *r*² = 0.047). Format as ind. **(f)** cue-*B* prelicking and DA↑^cueB^ emergence showed neither consistent temporal ordering nor detectable half-time covariance (two-sided sign test, *P* = 0.09; Pearson’s *P* = 0.57, *r*² = 0.026). Format as in **d.**

## Acknowledgements

We thank Michael Harris for preliminary visualizations of raw spike rasters; Ankit Choudhury for tyrosine hydroxylase immunostaining; Konstantin Bakhurin for training on electrode implantation; Robin Blazing for training on spike sorting; Isaac Weaver and Janani Sundararajan for assistance with building the behavioral arena; Vijay Namboodiri, Huijeong Jeong, Josh Dudman, Erin Calipari, Mark Harnett, Elias Issa, Rich Mooney, Steve Lisberger and Nicole Calakos for insightful feedback on the manuscript. This work was supported by Duke University Startup Funds, NIH grants RF1-MH117055 and DP2-MH1194025, and by the joint efforts of The Michael J. Fox Foundation for Parkinson’s Research (MJFF) and the Aligning Science Across Parkinson’s (ASAP) initiative. MJFF administers grant ASAP-020607 on behalf of ASAP and itself.

## Author Contributions

See **Table S2** for detailed author contributions. **Conceptualization:** S.C.V.B., M.R.T. **Methodology:** S.C.V.B., S.S.X.L., B.C.S., M.R.T. **Software:** S.C.V.B., M.R.T. **Validation:** S.C.V.B., H.Y., S.S.X.L., B.C.S., M.R.T. **Formal Analysis:** S.C.V.B., H.Y., M.R.T. **Investigation:** S.C.V.B., R.K.C., H.Y. **Resources:** S.S.X.L., B.C.S., M.R.T. **Data Curation:** S.C.V.B., R.K.C., H.Y. **Writing, Original Draft:** S.C.V.B., R.K.C., H.Y., M.R.T. **Writing, Review and Editing:** S.C.V.B., R.K.C., H.Y., S.S.X.L., B.C.S., M.R.T. **Visualization:** S.C.V.B., R.K.C., H.Y., M.R.T. **Supervision:** M.R.T. **Project Administration:** S.C.V.B., B.C.S., M.R.T. **Funding Acquisition:** M.R.T.

## IP Rights Notice

For the purpose of open access, the author has applied a CC-BY public copyright license to the Author Accepted Manuscript (AAM) version arising from this submission.

**Data and Software Availability** — All data and software are publicly available.

**Protocols:** https://doi.org/10.17504/protocols.io.j8nlk8ekdl5r/v1

**Software:** https://github.com/tadrosslab/VTA_GABA_paper and https://doi.org/10.5281/zenodo.10951255

## Datasets

- **Fig. 1, fig. S1-S2:** https://doi.org/10.5281/zenodo.10904059
- **Fig. 2, fig. S3:** https://doi.org/10.5281/zenodo.10903566 and https://doi.org/10.5281/zenodo.10908572
- **Fig. 3-4, fig. S4-S9:** supplementary data provided as excel files.

## Contact for Reagent and Resource Sharing

Further information and requests for resources and reagents should be directed to and will be fulfilled by the corresponding author Michael R. Tadross, MD, PhD.

## METHODS

### Mice

DAT-IRES-Cre (Jackson Labs 006660) mice were group housed by age and sex (max 5 per cage) in a standard temperature and humidity environment. For breeding, mice were housed under a normal 12-hr light/dark cycle and with food and water provided *ad libitum*. Experimental mice were transitioned to reverse-light-cycle and water-restriction conditions, as detailed below. All experiments involving animals were approved by the Duke Institutional Animal Care and Use Committee (IACUC), an AAALAC accredited program registered with both the USDA Public Health Service and the NIH Office of Animal Welfare Assurance, and conform to all relevant regulatory standards (Tadross protocols A160-17-06, A113-20-05, A091-23-04).

### Recombinant Adeno-associated Viral (rAAV) Vectors

All custom viral vectors were produced by the Duke Viral Vector Core or VectorBuilder, kept frozen at −80°C until use, then diluted to the desired titers using sterile hyperosmotic PBS and kept at 4°C for up to 4 weeks.

### Acute Brain Slice Electrophysiology

DAT-IRES-Cre mice (5 females, 3 males, 8-10 weeks) were anesthetized and stereotaxically injected with 400 nL of either AAV_rh10_-CAG-DIO-^+^HTP_GPI_-2A-dTomato-WPRE or AAV_rh10_-CAG-DIO-^dd^HTP_GPI_-2A-dTomato-WPRE (2 × 10^12^ VG/mL, 100 nL per site, two tracks with two depths per track: −3.2 mm AP, ±0.5 mm ML, −5.0/-4.5 mm DV) using a custom Narishige injector. After 3-5 weeks for expression, mice were deeply anesthetized with isoflurane and euthanized by decapitation. Coronal brain slices (300 µm) containing VTA were prepared by standard methods using a Vibratome (Leica, VT1200S), in ice-cold high sucrose cutting solution containing (in mM): 220 sucrose, 3 KCl, 1.25 NaH_2_PO4, 25 NaHCO_3_, 12 MgSO_4_, 10 glucose, and 0.2 CaCl_2_ bubbled with 95% O_2_ and 5% CO_2_. Slices were then placed into artificial cerebrospinal fluid (aCSF) containing (in mM): 120 NaCl, 3.3 KCl, 1.23 NaH_2_PO_4_, 1 MgSO_4_, 2 CaCl_2_, 25 NaHCO_3_, and 10 glucose at pH 7.3, previously saturated with 95% O_2_ and 5% CO_2_. Slices were incubated at 33°C for 40-60 min in bubbled aCSF and allowed to cool to room temperature (22-24°C) until recordings were initiated.

Recordings were performed on an Olympus BX51WI microscope, where slices were perfused with bubbled aCSF at 29-30°C with a 2 ml/min flow rate. To isolate GABA_A_ IPSCs, the external solution was supplemented with DNQX (20 µM, AMPA antagonist) and AP-V (50 µM, NMDA antagonist). Alternatively, to isolate AMPA-mediated EPSCs, aCSF was supplemented with picrotoxin (50 µM, GABA_A_R antagonist) and AP-V (50 µM). Finally, NMDA-mediated EPSCs were isolated with picrotoxin (50 µM) and DNQX (20 µM).

For voltage-clamp recording of GABA_A_R-IPSCs, the internal solution contained (in mM): 135 CsCl, 2 MgCl_2_, 0.5 EGTA, 10 HEPES, 4 MgATP, 0.5 NaGTP, 10 Na_2_-phosphocreatine, and 4 QX314, pH 7.3 with CsOH (290 mOsm). For voltage-clamp recordings of AMPAR or NMDAR EPSCs, the internal solution contained (in mM), 130 Cesium methanesulfonate, 2 MgCl_2_, 0.5 EGTA, 10 HEPES, 4 MgATP, 0. 5 NaGTP, 10 Na_2_-phosphocreatine, and 4 QX314, pH 7.3 with CsOH (290mOsm). For current-clamp, we used (in mM) 130 K-gluconate, 5 KCl, 2 MgCl_2_, 0.2 EGTA, 10 HEPES, 4 MgATP, 0.5 NaGTP, and 10 phosphocreatine, pH adjusted to 7.3 with KOH (290 mOsm). Internal solutions were used to fill glass recording pipettes (4-6 MΩ). The liquid junction potential, estimated to be 15.9 mV, was not corrected.

Whole-cell recordings were obtained with Multiclamp 700B, Digidata 1440A, pClamp 10.7 software (Molecular Devices). Signals were filtered at 10 kHz. A stimulating electrode was placed 60-100 μm from the recorded neuron. Evoked IPSCs or EPSCs were elicited by electrical stimuli of 0.3 ms duration and 150-300 μA (60-70% maximum responses), with a repetition interval of 15 sec. Our inclusion criteria required that cells maintain stable access and holding currents for at least 5 min. Series resistance was monitored using 5–10 mV hyperpolarizing steps interleaved with our stimuli, and cells discarded if series resistance changed more than 15%. The stored data signals were processed using Clampfit 10.7 (Axon Instruments).

### *In Vivo* Electrophysiology Experiments

Adult DAT-IRES-Cre mice (2 females, 6 males; 12-16 weeks old) were anesthetized and stereotaxically injected with 400 nL of either AAV_rh10_-CAG-DIO-^+^HTP_GPI_-2A-dTomato-WPRE, AAV_rh10_-CAG-DIO-^dd^HTP_GPI_-2A-dTomato-WPRE (2 × 10^12^ VG/mL), AAV_rh10_-CAG-CreON-W3SL-^+^HTP_GPI_-IRES-dTomato-Farnesylated, or AAV_rh10_-CAG-CreON-W3SL-^dd^HTP_GPI_-IRES-dTomato-Farnesylated (1 × 10^12^ VG/mL) (100nL per site, two tracks with two depths per track: −3.2 mm AP, ±0.5 mm ML, −5.0/-4.5 mm DV) with a custom Narishige injector. Mice were implanted with a single-drive movable micro-bundle electrode array (Innovative Neurophysiology, Inc.; 23 µm Tungsten Electrodes, 16 / bundle; 0.008” silver ground wire) above the left VTA (−3.2 mm AP, −0.5 mm ML, −4.0 mm DV). The silver ground wire was wrapped securely around two ground screws, one placed in the skull above the cerebellum and one above the right olfactory bulb. A unilateral metal cannula (P1Tech; C315GMN; cut to 13.5 mm) was implanted laterally adjacent to the electrode bundle (−3.2 mm AP, −1.3 mm ML, −4.0 mm DV). Mice were fitted with a plastic head bar adhered to the skull with OptiBond and dental cement. Mice were singly or pair housed post-surgery, in a 12-hr light/dark cycle, with food and water provided *ad libitum*. Pair-housed mice were outfitted with head hats that clip to specially designed head bars to prevent cannula or electrode damage from chewing by cage mates^28^.

Electrophysiology recordings and DART infusions were performed at least 3 weeks after surgery to allow for recombinant protein expression. The electrode bundle was manually advanced three times: (1) 208 µm at least one week after surgery, (2) another 208 µm one week later, and (3) 104 µm one week later. This placed the electrodes at −4.5 mm DV, at the top of the VTA. After a few days for recovery, electrophysiological recordings were made with an Intan RHD 16-channel headstage with accelerometer (C3335) attached to an Open Ephys Acquisition Board via an Intan RHD 1-ft ultra-thin SPI interface cable (C3211). Data were collected using the Open Ephys GUI^29^. Putative dopamine neurons were identified via their canonical features: tonic firing between 0 and 10 Hz, with bursting; wide biphasic or triphasic waveform; and large amplitude^22,23^. If no putative dopamine neurons were observed online, electrodes were advanced an additional 26-52 µm; this cycle was repeated until multiple channels with putative dopamine neurons were observed, at which point a recording was obtained.

DART ligands, stored as pure-compound aliquots, were freshly thawed on the day of use and dissolved in sterile artificial cerebrospinal fluid (aCSF) containing (in mM): 148 NaCl, 3 KCl, 1.4 CaCl_2_, 0.8 MgSO_4_, 0.8 Na_2_HPO_4_, 0.2 NaH_2_PO_4_. The final reagent solution contained 10 µM gabazine.7^DART.2^ + 1 µM alexa647.1^DART.2^. This solution was loaded into an internal cannula designed to project 0.5 - 1.5 mm from the guide cannula, with progressively longer internals used on successive infusions. Mice were head-fixed, the internal cannula inserted, and the Innovative Neurophysiology electrode bundle was connected to the Intan headstage. After obtaining a 15 min baseline recording, we infused 1.5 µL of DART reagent over 15 min (0.1 µL/min; Harvard Apparatus PhD Ultra pump; 5 µL Hamilton syringe), and continued the recording (120 min total). After completion of the recording, electrodes were advanced 26-52 µm^30^. Mice were given at least two weeks for recovery between recordings, which we have shown is sufficient to allow for complete HTP protein turnover^14^.

Spike sorting of the raw data was performed using SpyKING CIRCUS, an open-access software package allowing for semi-manual spike sorting on multichannel extracellular recordings^31^. Detection parameters included: spike threshold = 4; N_t (width of templates) = 2 or 3; peaks = positive. Filtering parameters used 250 Hz as the cutoff frequency for the Butterworth filter. All other parameters in the configuration file were standard as recommended by the SpyKING CIRCUS documentation. Only templates that matched all features of putative dopamine neurons and exhibited consistent spiking across the whole two-hour recording window were kept for analysis. All semi-manual spike sorting and template extraction were performed by S.C.V.B. for consistency.

Custom MATLAB code was used to extract the following metrics:

|  |  |  |  |
| --- | --- | --- | --- |
| <u>Tonic:</u> | *FR* | — | firing rate (spikes per second) |
|  | *mISI* | — | median interspike interval (ms) |
| <u>Pause:</u> | *fP* | — | fraction of $ISI > 2 \times mISI$ |
|  | *PL* | — | pause length, normalized (mean pause *ISI* divided by *mISI*, unitless) |
| <u>Burst:</u> | *fB* | — | fraction of spikes fired in bursts |
|  | *SPB* | — | mean spikes per burst (unitless) |

All metrics were calculated within 15-min windows. Pause metrics compared each *ISI* to the *mISI* over the same 15-min window. Bursts were defined as a sequence of 3-10 spikes in which the first *ISI* < 80 ms and subsequent *ISI* < 160 ms^22,23^. Changes in a given metric, *m*, were analyzed by comparing the 15-min baseline (*m*_pre_) to a 15-min sliding window (*m*_post_) according to: *Δm* = (*m*_post_ – *m*_pre_) / (*m*_post_ + *m*_pre_). We plotted the time course of *Δm* (as a function of the sliding-window time) and analyzed the steady-state *Δm* (1-hr post-gabazine^DART^) using a two-sided permutation test^32^. Correlations between metrics were analyzed with a Pearson’s test.

### Behavior Experiments without Photometry

Adult DAT-IRES-Cre mice (20 male, 17 female; 12-16 weeks old) were anesthetized and stereotaxically injected in the VTA with 400 nL of AAV_rh10_-CAG-DIO-^+^HTP_GPI_-2A-dTomato-WPRE or AAV_rh10_-CAG-DIO-^dd^HTP_GPI_-2A-dTomato-WPRE (2 × 10^12^ VG/mL, 100 nL per site, two tracks with two depths per track: −3.2 mm AP, ±0.5 mm ML, −5.0/-4.5 mm DV) with a custom Narishige injector. Mice were implanted with a bilateral metal cannula above the VTA (P1Tech; C235G-1.0; cut to 4 mm with 1.0 mm spacing), which was lowered slowly to −3.75 mm. Mice were fitted with a plastic head bar adhered to the skull with OptiBond and dental cement, enabling head fixation. Mice were singly or pair housed post-surgery, in a 12-hr reverse light/dark cycle, with food and water provided *ad libitum*. Pair-housed mice were outfitted with head hats that clip to specially designed head bars^28^ to prevent cannula damage from chewing by cage mates.

Mice were given a minimum of 9 days post-surgery for recovery and acclimation to the reverse light cycle. For the subsequent 3 days, mice were habituated to head-fixation and water restriction. Water was limited to 50-60 µL per gram of the mouse’s baseline weight per day, while dry food was provided *ad libitum*. The water restriction goal was 85% starting body weight; additional supplementary water was provided if mice dropped below 77% original body weight or did not pass a daily qualitative health assessment. Only 1 mouse was excluded for issues with water restriction health.

During behavioral sessions, mice were head-fixed^28^ on a round plastic treadmill (Delvie’s Plastics, 8” plexiglass disk covered with silicone rubber) attached to a rotary encoder to collect rotation data (U.S. Digital H5-100-NE-S). Cue tones were played through a Z50 speaker, lick detection was collected with an infrared beam, and sucrose rewards were delivered via a Lee Company solenoid (LHDA1233315H HDI-PTD-Saline-12V-30PSI). Before each behavioral session, the solenoid opening duration was calibrated to deliver 5 μL of freshly prepared 10% sucrose solution per activation. A custom MATLAB script controlled the behavioral sessions and data collection via a National Instruments card (NI USB-6351 X Series DAQ). Behavior sessions lasted 1 hr per day for 12 consecutive days and were performed during the dark portion of the mouse’s circadian cycle. The order in which mice performed the task was pseudorandomly counterbalanced through day 11. On day 12, mice were tested in the same order as on day 11 to maintain a consistent interval from gabazine^DART^ infusion to the second testing session.

During training sessions (days 1-10), mice were conditioned to associate cue *A* (2.5 kHz tone, 1.5 sec) with a 5 µL 10% sucrose-water reward. Conditioning trials were randomly interleaved with silent trials (with neither cue nor reward), enabling consistency in the trial-structure and reward-delivery quantities throughout training and testing sessions. On the final day of training (day 10) we replaced 5 of the silent trials with probe trials in which an unfamiliar cue *B* (11 kHz tone, 1.5 sec) was presented but unrewarded. Thereafter, on day 11, we infused 10 µM gabazine.7^DART.2^ + 1 µM alexa647.1^DART.2^ dissolved in sterile aCSF; 0.6 - 0.8 µL was infused per hemisphere at a rate of 0.1 µL/min (Harvard Apparatus PhD Ultra pump using 5 µL Hamilton syringes). Following a 2 hr rest, mice resumed the original training rules for 15 min. Thereafter the rules changed: cue *A* was now unrewarded (extinction trials) interleaved with cue *B* rewarded (conditioning trials). These rules continued on day 12. Throughout the assay, mice completed 200–300 total trials daily (half cue-*A*; half silent or cue-*B*). Licks were allowed during the 1.5 sec tone (anticipatory ‘prelicks’) and the subsequent 2 sec period (retrieval licks). The inter-trial interval (ITI) was random 3 - 13 sec (from the end of the retrieval period to the start of the next cue). Licks during the ITI restarted the interval timer (without rerandomizing its duration) to discourage nonspecific licking. Time penalties were never imposed for licking during a cue or retrieval period (regardless of whether the cue was rewarded or unrewarded). Prelicking (during the 1.5 sec cue) was our primary learning measure, which we quantify as the number times that the infrared beam was broken per second (Hz) during the cue.

Following the session on day 12, all mice were perfused for histological visualization of tracer^DART^ capture. No mice were excluded based on histology. Our behavioral inclusion criteria required that mice exhibit mean cue-*A* prelicking greater than 1 Hz on the 10^th^ training session (this was satisfied by 25/36 mice), and cue-*B* probe-trial prelicking less than 30% of responses to cue-*A* (satisfied by 24/25 mice). Thus, the behavior-only cohort comprised 24 mice that met the behavioral inclusion criteria (12 ^dd^HTP, 12 ^+^HTP). Behavioral analyses pooling this cohort with the photometry cohort comprised 39 mice in total (20 controls and 19 ^+^HTP/gabazine^DART^ mice). The behavioral experimenter was blinded to virus condition in half of the experimental cohorts. All statistical comparisons used two-sided permutation tests^32^.

### Behavior Combined with Photometry

In a later photometry cohort, all mice expressed ^+^HTP and were assigned after pre-infusion signal-quality assessment to receive either 10 µM gabazine.7^DART.2^ + 1 µM alexa647.1^DART.2^ or 10 µM blank^DART^ + 1 µM alexa647^DART^. This enabled counterbalanced treatment allocation after assessment of baseline signal fidelity. The original ^dd^HTP/gabazine^DART^ (*n* = 12) and the subsequent ^+^HTP/blank^DART^ (*n* = 8) controls did not differ in any behavioral metric (Δlocomotion: *P*_unadj_ ≥ 0.096, *P*_Holm_ ≥ 0.67; Δlick: *P*_unadj_ ≥ 0.072, *P*_Holm_ ≥ 0.50; and *t*_50%_: *P*_unadj_ ≥ 0.11, *P*_Holm_ ≥ 0.22). Similarly ^+^HTP/gabazine^DART^ from the original (*n* = 12) and photometry (*n* = 7) cohorts did not differ in any behavioral metric (Δlocomotion: *P*_unadj_ ≥ 0.54, *P*_Holm_ = 1; Δlick: *P*_unadj_ ≥ 0.23, *P*_Holm_ = 1; and *t*_50%_: *P*_unadj_ ≥ 0.22, *P*_Holm_ ≥ 0.44). We therefore pooled behavioral data where indicated.

Adult DAT-IRES-cre mice (15 male, 12 female) were stereotaxically injected in the VTA with AAV_rh10_-CAG-DIO-^+^HTP_GPI_-2A-dTomato-WPRE (2 × 10^12^ VG/mL, 100 nL per site, two tracks with two depths per track: −3.2 mm AP, ±0.5 mm ML, −5.0/-4.5 mm DV) and in the left lateral NAc with AAV9-hSYN-GRAB_DA3m_ (1 × 10^13^ VG/mL, 100 nL per site, one track with two depths: +1.6 mm AP, −1.5 mm ML, −4.0/-4.5 mm DV). Mice were implanted with a bilateral cannula above the VTA (P1Tech; C235G-1.0; cut to 4 mm with a 1.0 mm spacing) lowered to −3.75 mm, and an optic fiber (Doric Lenses, MFC_400/430-0.66_5mm_MF1.25_FLT) targeting the lateral NAc −4.0 mm DV. Headcaps, reverse-light cycle housing, and behavioral timeline were as noted previously.

GRAB_DA3m_ fluorescence was recorded using a Tucker-Davis Technologies RZ10X processor and Synapse software. Excitation light was delivered through a mono fiber-optic patch cord at 415 nm and 465 nm, adjusted to 20 μW and 25 μW, respectively, at the patch-cord output. Photometry recording began before each behavioral session and continued until the session ended. Behavioral and photometry data were synchronized by recording the solenoid-command signal through an analog input on the RZ10X. Analyses used only the 465 nm channel.

We introduced two additional kinds of probe trials to facilitate photometry calibration to relevant pre-infusion values. To measure DA dips to unexpected omission before drug infusion, we introduced rare *A*→omit probes on days 7-10 (three per day), which became the pre-infusion values used to calculate ΔDA↓^omit^. Similarly, we measured DA elevations to unexpected reward by introducing rare uncued rewards on days 7-10 (three per day), which became the pre-infusion values used to calculate ΔDA↑^reward*B*^.

As before, we established that mice could discriminate cues by replacing 5 of the silent trials on day 10 with probe trials in which an unfamiliar cue *B* (11 kHz tone, 1.5 sec) was presented but unrewarded. Mice failing behavioral inclusion criteria were excluded (4 for insufficient *A*-prelicking; 1 for insufficient *B*-discrimination), and one mouse was excluded due to histological evidence of a large VTA lesion. Photometry signal fidelity was quantified using two normalized metrics: mean cue-*A*-evoked *ΔF/F*_0_ across days 6-10 relative to its standard deviation over trials, and probe-omission dip magnitude relative to its standard deviation over trials. The minimum of these two values was used as a conservative signal-fidelity measure, and mice were required to exceed a predefined threshold (fidelity > 1) to ensure adequate signal quality. Of the 21 mice that passed behavioral and histological inclusion criteria, 15 mice (7 ^+^HTP/gabazine^DART^, 8 controls) passed signal-quality criteria and were included in the final dataset.

### Photometry Waveform Analysis

We first computed *DA =* 100 *×* (*F – F*₀) */ F*₀ where *F* is the instantaneous GRAB_DA3m_ fluorescence, and *F*₀ is the baseline fluorescence.

*F*₀ was estimated using an iterative moving-median procedure. A 30-s moving median was first calculated from the fluorescence trace, yielding *F*₀₁, a preliminary estimate of *F*₀. We then calculated *DA*₀₁ = 100 *×* (*F* − *F*₀₁) / *F*₀₁. Samples whose absolute *DA*₀₁ exceeded the magnitude of the first percentile of the *DA*₀₁ distribution were temporarily masked in the original fluorescence trace before recalculating the moving median, yielding *F*₀₂. At each iteration, the exclusion threshold was recalculated from the updated *DA* distribution. Missing values in the resulting baseline estimate were filled by linear interpolation, with edge values filled using the nearest valid estimate. This procedure was repeated for 10 iterations, progressively reducing the influence of large upward-and downward-going transients on the baseline estimate. The final moving-median estimate was designated *F*₀.

Because photometry does not resolve electrophysiologically defined pauses and bursts, the continuous DA waveform was separated into downward-going and upward-going components by half-wave rectification. For the upward-going component, DA↑(t), samples below zero were set to zero. For the downward-going component, samples above zero were set to zero and the remaining negative values were sign-inverted to yield DA↓(t). Thus, DA↓(t) and DA↑(t) were both non-negative, whereas the arrows indicate the original direction of fluorescence deflection. Samples set to zero were retained in the continuous traces so that the mean rectified signal within an epoch reflected both the amplitude and prevalence of deflections in the analyzed direction.

DA↓(t) and DA↑(t) were analyzed separately within seven predefined behavioral epochs: intertrial intervals (ITI), cue *A* before the rule flip (preflipCue*A*), reward following cue *A* before the rule flip (reward*A*), cue *A* after the rule flip (cue*A*), reward omission after cue *A* (omit), cue *B* after the rule flip (cue*B*), and reward following cue *B* (reward*B*). Cue epochs comprised the 1.5-s cue period. Outcome epochs comprised the subsequent 2-s retrieval or omission period. ITI epochs extended from the last lick, which restarted the ITI timer, to the onset of the next cue. For each trial, the mean rectified signal within the corresponding epoch was calculated while retaining all zero-valued samples introduced by rectification. Trial-level values were assigned to their elapsed times within the assay and placed on a common minutes-scale time axis for visualization.

Pre-infusion reference periods were defined separately for each epoch. DA↓^ITI^ and DA↑^ITI^ used ITIs from training days 7–10. PreflipCue*A* and post-flip cue*A* signals used cue-*A* trials from training days 7–10. Reward*A* signals used rewards following cue *A* on training days 7–10. Omission signals used three probe omissions per day on training days 7–10. Cue*B* signals used cue-*A* trials from training days 7–10, because cue *B* was novel before the rule flip. Reward*B* signals used uncued-reward probes delivered on training days 7–10. For preflipCue*A* and reward*A*, the post-infusion period was the 15-min interval preceding the rule flip on day 11. For cue*A*, omit, cue*B*, and reward*B*, the post-infusion period comprised the post-flip portions of days 11 and 12. For ITI analyses, the post-infusion period comprised the full 120-min assay spanning the pre-flip interval, the post-flip portion of day 11, and day 12.

For each signal direction and behavioral epoch, normalized within-mouse change was calculated as Δ = (post − pre) / (post + pre), where pre and post were the corresponding mean pre-and post-infusion values. For DA↓ measures, negative Δ values indicate shallower or less prevalent downward-going deflections after infusion. For DA↑ measures, negative Δ values indicate smaller or less prevalent upward-going deflections after infusion. Unless otherwise specified, full-period Δ values used the complete post-infusion period defined above.

For cumulative analyses, the pre-infusion mean was held constant while the post-infusion term was calculated as the cumulative mean from the beginning of the relevant post-infusion period to the plotted time. For DA↓^ITI^, the first point was plotted at 15 min and used the mean from 0–15 min after infusion; the value at 60 min used the mean from 0–60 min; and the full-period value used the mean from 0–120 min. For DA↓^omit^, cumulative averaging began at the rule flip and included all omission trials through the plotted time. Early ΔDA↓omit used the first four omissions after the rule flip.

For visualization of photometry time courses, all values from each treatment group were multiplied by a single group-specific constant so that the day 7–10 pre-infusion mean for that group equaled the geometric mean of the two group baselines. This group-level rescaling preserved relative within-group variability and did not normalize individual mice. All statistical analyses were performed on unscaled within-mouse values. Raw values are provided as supplementary data. Half-times were estimated separately for cue-*A* prelicking, DA↑^cue*A*^, DA↓^omit^, cue-*B* prelicking, DA↑^cue*B*^, and DA↑^rewardB^. For each mouse, the post-flip trajectory was fit with a logistic function. The half-time, *t*_50%_, was defined as the time at which the fitted trajectory reached the midpoint between its fitted initial and asymptotic values. For decreasing trajectories, this represented half-dissipation; for increasing trajectories, it represented half-emergence.

Treatment effects were assessed using two-sided permutation tests. Downward-going dopamine signals and upward-going dopamine signals were treated as separate families, each comprising the seven predefined behavioral epochs, and *P* values within each family were adjusted using the Holm procedure. Associations between continuous measures were assessed using two-sided Pearson correlations. Temporal ordering between paired half-times was assessed using exact two-sided sign tests relative to the line of equal half-times. Multivariable linear models tested whether DA↓^omit^ half-time accounted for behavioral half-extinction time after adjustment for treatment and whether this relationship differed by treatment. The slope relating behavioral half-extinction time to DA↓^omit^ half-dissipation time was additionally tested against a null slope of 1. A slope greater than 1 indicated that the delay between DA↓^omit^ dissipation and behavioral extinction increased as DA↓^omit^ persisted longer.

### Locomotion analysis

Treadmill rotation was recorded continuously using a quadrature rotary encoder (U.S. Digital H5-100-NE-S) coupled directly to the circular treadmill and acquired through the NI USB-6351 data-acquisition card. Encoder displacement was converted to wheel rotations per min (RPM). Locomotion was averaged separately within the same seven behavioral epochs used for the photometry analyses: ITI, preflipCue*A*, reward*A*, cue*A*, omit, cue*B* and reward*B*. Cue epochs comprised the 1.5-s cue period, outcome epochs comprised the subsequent 2-s retrieval or omission period, and ITI epochs extended from the final lick to the onset of the next cue. For all seven locomotion metrics, the common pre-infusion reference was mean ITI locomotion across training days 7–10. Normalized within-mouse changes were calculated as Δ = (post − pre) / (post + pre), where post was the mean locomotion during the corresponding post-infusion epoch. Treatment effects across the seven locomotion epochs were treated as one family and adjusted using the Holm procedure. The display-only group rescaling described for photometry was applied analogously to the locomotion time courses; statistical analyses used unscaled within-mouse values.

### Logistic fitting

Half-times were estimated separately for prelick*A*, DA↑^cue*A*^, DA↓^omit^, prelick*B*, DA↑^cue*B*^ and DA↑^reward*B*^. For each mouse and measure, the post-infusion waveform was normalized to the magnitude of its pre-infusion reference, *R*(t) = *post*(*t*) / |mean(*pre*)|, using the variable-specific reference periods defined above. Normalized trajectories were smoothed with a five-minute moving mean using MATLAB smoothdata. The post-flip time axis was defined so that the first observation occurred at *t* = 0. Trajectories were fitted by robust nonlinear least squares. We defined *Q*(*t*) = (exp(*h*/*τ*) – 1) / (exp(*h*/*τ*) + exp(*t*/*τ*) – 2), where *h* is the transition half-time and *τ* controls transition steepness. Because *Q*(0) = 1, *Q*(*h*) = 0.5, and *Q*(*t*)→0 as *t*→∞, decreasing trajectories were fitted as *R*(*t*) = *z* + (1 - *z*) *Q*(*t*), where *z* is the fitted lower asymptote. Cue-*A* prelicking was constrained to *z* = 0; for DA↑^cue*A*^, DA↓^omit^ and DA↑^reward*B*^, *z* was constrained to 0 – 0.05. Increasing cue-*B* prelicking and DA↑^cue*B*^ trajectories were fitted as *R*(*t*) = *u*(1 - *Q*(*t*)), where the fitted upper asymptote, *u*, was constrained to 0.2–1. For decreasing trajectories, *h* was constrained to 1–100 min; for increasing trajectories, it was constrained to 10–100 min. For all fits, *τ* was constrained to 3–30 min. The fitted value of *h*, denoted *t*_50%_, therefore represented half-dissipation for decreasing trajectories and half-emergence for increasing trajectories.

### Histology

Mice were deeply anesthetized with isoflurane. Electrodes were briefly connected to a 9V battery (1 sec) to mark electrode positions. Thereafter, mice were fixed by transcardial perfusion of 15 mL PBS followed by 50 mL ice-cold 4% paraformaldehyde (PFA) in 0.1M PB, pH 7.4. Brains were excised from the skull, post-fixed in 50 mL of 4% PFA at 4°C overnight, then washed three times with PBS. Brains were embedded in 5% agarose and sliced along the coronal axis at 50 µm (Leica, VT1200S).

For tyrosine hydroxylase immunostaining, sections were washed in PBS before a 2 hr incubation in a blocking solution consisting of 5% goat serum, 3% bovine serum albumin, and 0.3% Trition X-100. Sections were then transferred to a half block solution containing 1:1000 rabbit anti-TH (PelFreez, P40101) overnight at 4°C with agitation, and then washed in 0.1M PBS containing 0.1% tween before a 4 hr incubation in a half block solution containing 1:1000 goat anti-rabbit 488 (Invitrogen, A11008). Finally, sections were washed in PBS containing tween, then PBS alone prior to mounting on glass slides.

Sections were mounted onto glass slides (VWR 48311-703) and coverslipped with Vectashield mounting medium (Vector Labs, H-1400 or H-1800). Fluorescent images (DAPI, FITC, TRITC, Cy5) were collected at 10X magnification with an Olympus VS200 slide scanner.

Cell counts were obtained using ilastik^33^. Pixel Classification was used to predict cell versus not-cell (background), then Object Classification was used on these pixel predictions to label cells as red (dTomato), green (TH+), or red+green (both). Object identities were used to calculate the number of cells identified in each label class across all sections from one brain.

### Use of generative AI

Large language models were used to assist with language editing and critical review of manuscript text. All analyses, interpretations and final wording were independently evaluated and approved by the authors, who take full responsibility for the content.

---

**Table S1:**
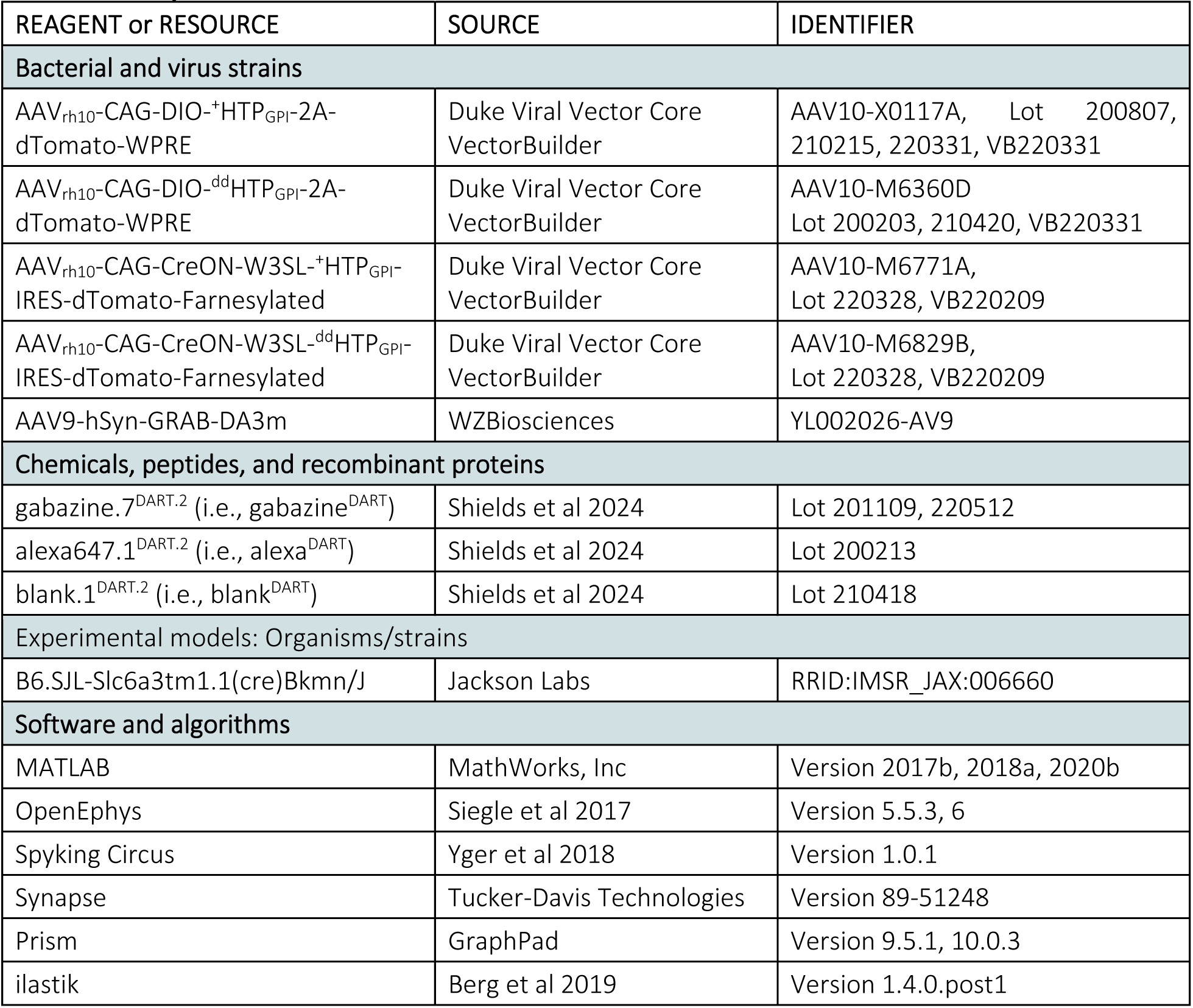
Key Resources.

| REAGENT or RESOURCE | SOURCE | IDENTIFIER |
| --- | --- | --- |
| **Bacterial and virus strains** |  |  |
| AAV<sub>rh10</sub>-CAG-DIO-<sup>+</sup>HTP<sub>GPI</sub>-2A-dTomato-WPRE | Duke Viral Vector Core VectorBuilder | AAV10-X0117A, Lot 200807, 210215, 220331, VB220331 |
| AAV<sub>rh10</sub>-CAG-DIO-<sup>dd</sup>HTP<sub>GPI</sub>-2A-dTomato-WPRE | Duke Viral Vector Core VectorBuilder | AAV10-M6360D<br>Lot 200203, 210420, VB220331 |
| AAV<sub>rh10</sub>-CAG-CreON-W3SL-<sup>+</sup>HTP<sub>GPI</sub>-IRES-dTomato-Farnesylated | Duke Viral Vector Core VectorBuilder | AAV10-M6771A,<br>Lot 220328, VB220209 |
| AAV<sub>rh10</sub>-CAG-CreON-W3SL-<sup>dd</sup>HTP<sub>GPI</sub>-IRES-dTomato-Farnesylated | Duke Viral Vector Core VectorBuilder | AAV10-M6829B,<br>Lot 220328, VB220209 |
| AAV9-hSyn-GRAB-DA3m | WZBiosciences | YL002026-AV9 |
| **Chemicals, peptides, and recombinant proteins** |  |  |
| gabazine.7<sup>DART.2</sup> (i.e., gabazine<sup>DART</sup>) | Shields et al 2024 | Lot 201109, 220512 |
| alexa647.1<sup>DART.2</sup> (i.e., alexa<sup>DART</sup>) | Shields et al 2024 | Lot 200213 |
| blank.1<sup>DART.2</sup> (i.e., blank<sup>DART</sup>) | Shields et al 2024 | Lot 210418 |
| **Experimental models: Organisms/strains** |  |  |
| B6.SJL-Slc6a3tm1.1(cre)Bkmn/J | Jackson Labs | RRID:IMSR_JAX:006660 |
| **Software and algorithms** |  |  |
| MATLAB | MathWorks, Inc | Version 2017b, 2018a, 2020b |
| OpenEphys | Siegle et al 2017 | Version 5.5.3, 6 |
| Spyking Circus | Yger et al 2018 | Version 1.0.1 |
| Synapse | Tucker-Davis Technologies | Version 89-51248 |
| Prism | GraphPad | Version 9.5.1, 10.0.3 |
| ilastik | Berg et al 2019 | Version 1.4.0.post1 |

**Table S2.**
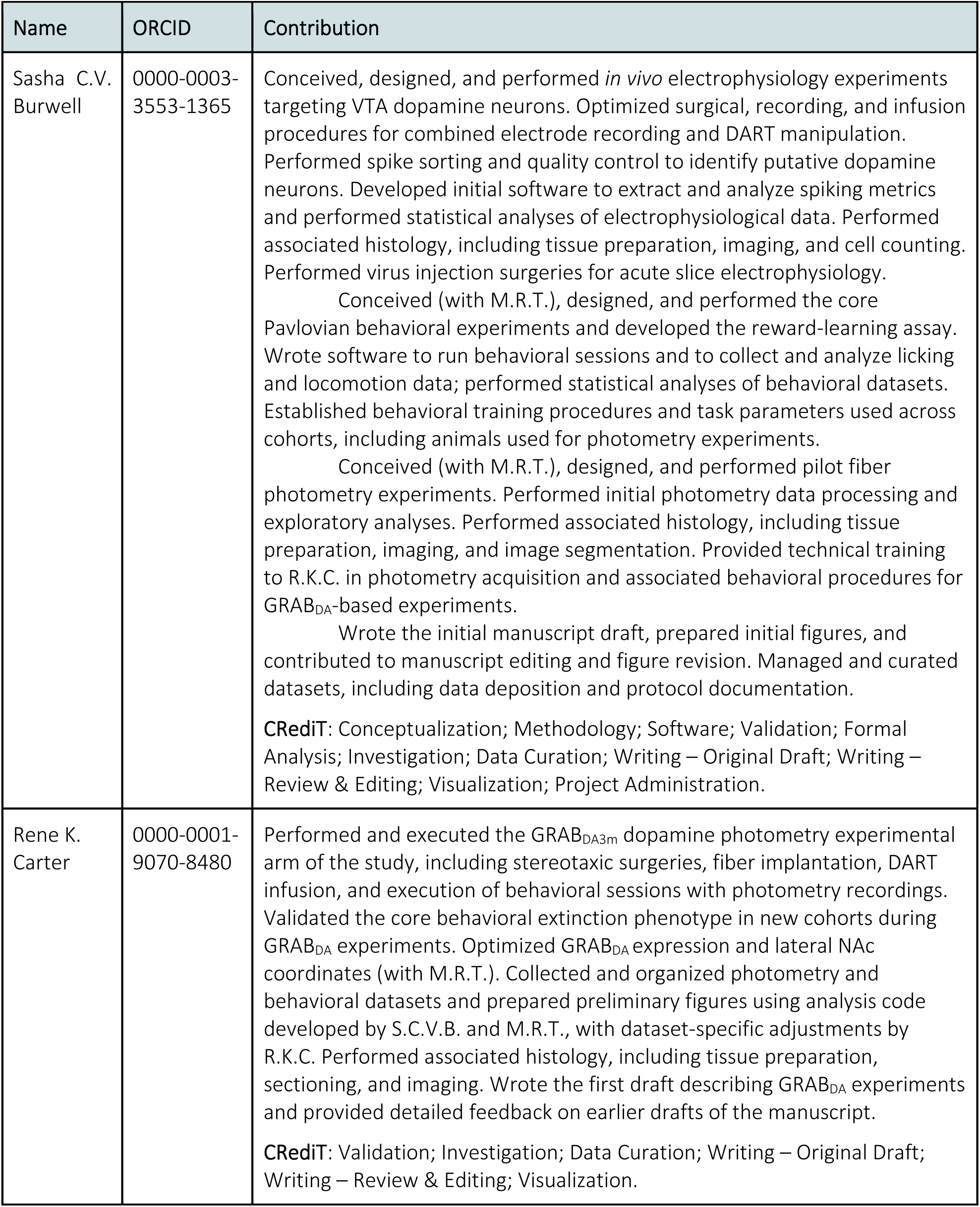
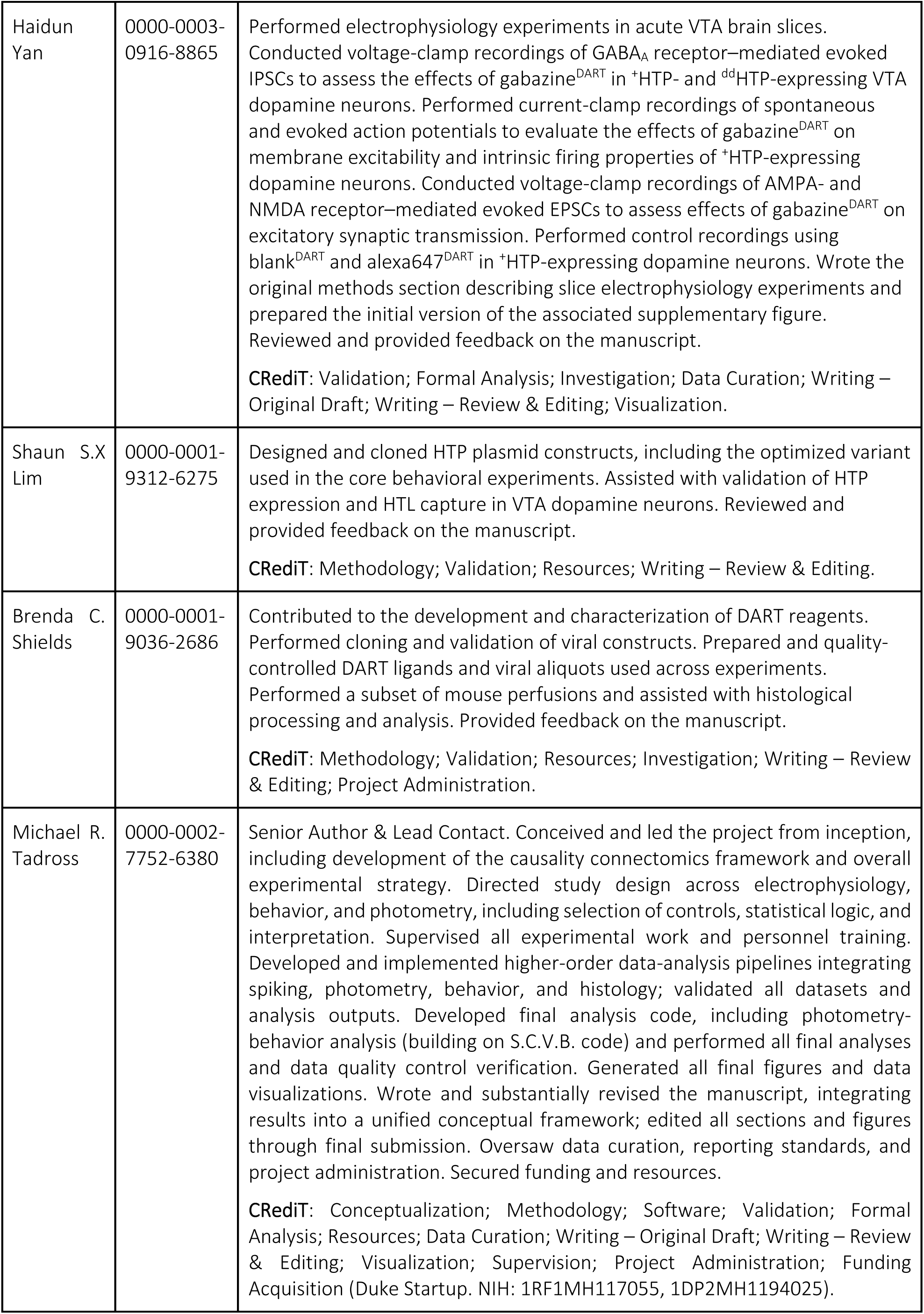
Detailed Author Contributions.

